# A quantitative survey of the blueberry (*Vaccinium* spp.) nectar microbiome: variation between cultivars, locations, and farm management approaches

**DOI:** 10.1101/2023.09.11.556904

**Authors:** Caitlin C. Rering, Arthur B. Rudolph, Qin-Bao Li, Quentin D. Read, Patricio R. Muñoz, John J. Ternest, Charles T. Hunter

**Author notes:** Corresponding author:**, 1700 SW 23^rd^ Dr, Gainesville, FL 32608;.

## Abstract

Microbes in floral nectar can impact both their host plants and floral visitors, yet little is known about the nectar microbiome of most pollinator-dependent crops. In this study, we examined the abundance and composition of the fungi and bacteria inhabiting *Vaccinium* spp. nectar, as well as nectar volume and sugar concentrations, hypothesizing that nectar traits and microbial communities would vary between plants. We compared wild *V. myrsinites* with two field-grown *V. corymbosum* cultivars collected from two organic and two conventional farms. Differences in nectar traits and microbiomes were identified between *V. corymbosum* cultivars but not *Vaccinium* species. The microbiome of cultivated plants also varied greatly between farms, whereas management regime had only subtle effects, with higher fungal populations detected under organic management. Nectars were hexose-dominant, and sugars were depleted in nectar with higher cell densities. Bacteria were more common than fungi in blueberry nectar, although both were frequently detected and co-occurred more often than would be predicted by chance. ‘Cosmopolitan’ blueberry nectar microbes that were isolated in all plants, including *Rosenbergiella* sp. and *Symmetrospora symmetrica*, were identified. This study provides the first systematic report of the blueberry nectar microbiome, which may have important implications for pollinator and crop health.

**One-sentence summary:** Parallel analysis of blueberry crops and a wild relative offers insight into the impacts of management and domestication on the nectar microbiome

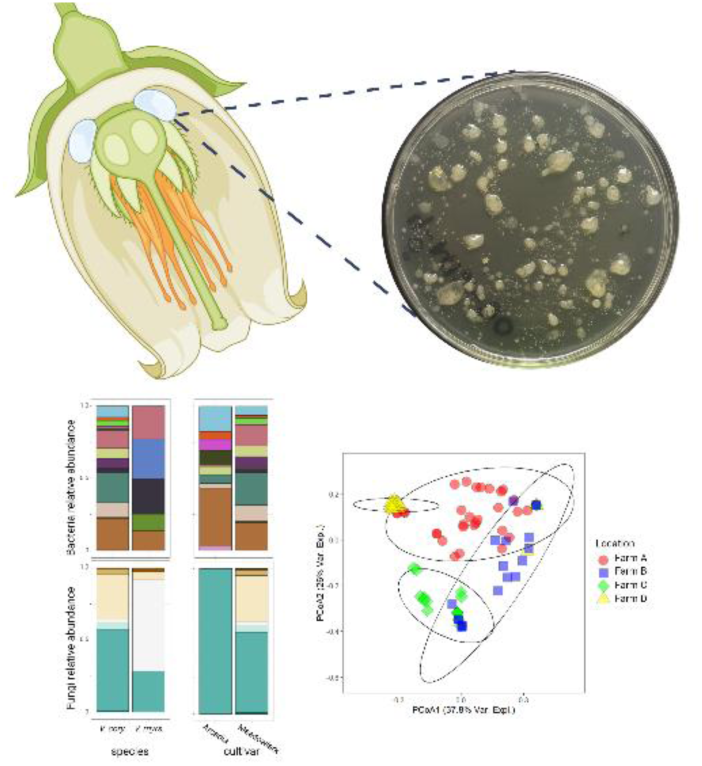

## Introduction

Nectar, the major reward used to attract pollinators to flowers, is frequently colonized by microbes (Herrera *et al*. 2009; Álvarez-Pérez, Herrera and de Vega 2012; Álvarez-Pérez and Herrera 2013; de Vega *et al*. 2021). Nectar microbes can have significant impacts on both their host plant and visiting insects (reviewed in Vannette 2020). For example, microbes change the chemical composition of nectar, thereby influencing pollinator health (Pozo *et al*. 2020; Pozo *et al*. 2021; reviewed in Martin, Schaeffer and Fukami 2022) and preference for flowers (Herrera, Pozo and Medrano 2013; Vannette *et al*. 2013; Schaeffer et al 2014; Yang *et al*. 2019; Colda *et al*. 2021). Microbial modifications to nectar include metabolizing nutrients like sugars and amino acids (Canto and Herrera 2012; Vannette and Fukami 2018), metabolizing phytochemicals (Vannette and Fukami 2016), and producing volatile (Rering *et al*. 2018; Sobhy *et al*. 2018; 2019; Cusumano *et al*. 2022) and non-volatile metabolites (Schaeffer *et al*. 2019), all of which can impact pollinator affinity to nectar. In turn, microbial modulations to floral attractiveness can affect plant reproduction by increasing or decreasing visitation to flowers (Vannette *et al*. 2013; Schaeffer and Irwin 2014; Yang *et al*. 2019; de Vega et al 2022). Nectar– and flower-inhabiting microbes can also have direct impacts on plant health either as pathogens (Bubán, Orosz-Kovács, Farkas 2003; Ngugi and Scherm 2006) or as protective species that can prevent or reduce disease (Pusey, Stockwell and Mazzola 2009; Chacón *et al*. 2022; Crowley-Gall *et al*. 2022).

The floral nectar microbiome is usually comprised of so-called nectar specialists, microbes that are isolated from the nectar of many plants but rarely elsewhere, and a subset of phyllosphere– and insect-associated species that can withstand the nectar niche (Herrera *et al*. 2010; Pozo, Lachance, Herrera 2012; reviewed in Vannette 2020). Nectar is a challenging environment for microbial growth, characterized by extreme osmotic pressure due to high sugar concentrations, antimicrobial plant defense chemistry (e.g., phytochemicals and proteins), limited nitrogenic resources, and intense UV radiation (Heil 2011). Even so, nectar is frequently colonized by microbes that can reach high cell densities (Herrera *et al*. 2009; Álvarez-Pérez and Herrera 2013; de Vega *et al*. 2021; Vannette *et al*. 2021). As flowers age, microbial incidence and abundance tends to increase as visitors, wind, and rain vector microbes to flowers (Herrera *et al*. 2010; Aizenberg-Gershtein, Izhaki, Halpern 2013; Vannette and Fukami 2017; von Arx *et al*. 2019; Morris *et al*. 2020). Insect dispersal is particularly important in shaping communities, with differing insects having distinct impacts on the nectar microbiome (Morris *et al*. 2020; Zemenick, Vannette, Rosenheim 2021; de Vega *et al*. 2021). Plant species and location are also important factors (Fridman *et al*. 2012; Jacquemyn *et al*. 2013a; Aizenberg-Gershtein, Izhaki, Halpern 2017; Vannette *et al*. 2021; Schaeffer *et al*. 2021) and in crops, the application of fungicides have been shown to impact nectar communities (Schaeffer *et al*. 2017a; 2021; 2023; Wei *et al*. 2021).

In the last 20 years, research describing the nectar microbiome has intensified: flowers across the globe have been surveyed, from the tropics (Canto and Herrera 2012) to the Artic (Niu et al 2023). Although most studies have focused on natural habitats, some have also examined the nectar microbiome within agroecosystems, predominantly in orchards. Very few studies have examined the nectar communities in non-orchard perennial crops (but see research in coffee nectar, Vannette *et al*. 2017). Here, we report the first study of blueberry (*Vaccinium* spp.) nectar microbial communities. Blueberry is a highly nutritious pollinator-dependent crop that generates over $1 billion annually in the US (U.S. National Agricultural Statistics Service 2021). To date, only one study has reported microbes inhabiting the blueberry anthosphere (Chacón *et al*. 2022), with the aim of isolating bacteria with biocontrol potential from whole blueberry flowers and fruit, rather than providing a systematic survey of microbial incidence and abundance in nectar.

We studied two commercial highbush blueberry cultivars (*Vaccinium corymbosum*), one high-yielding and the other suffering from low yields. Both cultivars were collected from organic and conventional farms. We compared cultivated *V. corymbosum* with a wild blueberry species native to the southeastern USA, *Vaccinium myrsinites* or shiny blueberry (Wunderlin *et al*. 2023). Because domestication of blueberries has modified nectar and pollen chemistry with potential impact on pollinator attraction (Egan *et al*. 2018), and these factors are known to affect the nectar microbiome, we predicted that nectar from wild and cultivated plants would host differing microbial communities. Although a few studies have explored the floral microbiome in wild relatives of domesticated crops (wild strawberry, Wei and Ashman 2018; wild *Brassica rapa,* Russell and McFrederick 2022), this is the first concurrent comparison of crops and wild relatives present in similar temporal and geographic locales, providing insight into the effects of domestication on crop microbiomes. Similarly, blueberry cultivars differ in reward chemistry and insect visitation, leading to unique communities. Only one study has examined differences in floral communities between crop cultivars, finding the floral microbiome varied between three cultivated strawberry lines (Wei *et al*. 2022).

We had three goals in this study: first, to characterize the microbial community present in blueberry nectar; second, to evaluate whether nectar microbial communities differed between blueberry species, cultivars, organic vs. conventional management, and location; third, to evaluate nectar volume and chemistry offered by each plant and examine what, if any, impact microbes had on nectar sugars. We hypothesized that blueberry nectar would contain a “core” microbiome that included nectar specialists, but that differences among cultivars, plant species, management, and location would be evident.

## Materials and Methods

### Collection sites and species

Flowers were collected from four farms and two nature preserves within a 130 mi range of North and Central Florida (see Fig. S1 for sampling sites). Flowers were collected when plants were at peak bloom for a given location, between 9:00 AM and 2:30 PM from January 30^th^ – March 3^rd^, 2020. Two cultivars of highbush blueberry (*Vaccinium corymbosum*; Ericaceae), Meadowlark and Arcadia, were investigated in this study. Florida blueberry farmers report Meadowlark as low-yielding (Phillips 2019a) and Arcadia as high-yielding (Phillips 2019b). Both *V. corymbosum* cultivars were sampled from four farms: two organic and two conventional. Shiny blueberry (*Vaccinium myrsinites*; Ericaceae) flowers were collected from Austin Cary and Ross Prairie State Forests.

*V. corymbosum* blueberry flower samples were collected from individual plants distributed throughout the field. *V. myrsinites* blueberry propagates clonally, often forming dense stands near one another (5‒10 m^2^). For each forest surveyed, flowers were collected from 2‒3 areas, each separated by several hundred meters. *V. myrsinites* flowers were collected from individual shoots throughout an area. For both *Vaccinium* species, only open, intact flowers with no signs of senescence, nectar robbing, or herbivory were collected. Flowers were cut from the plant and placed in sterile 2 mL centrifuge tubes. Tubes were loaded into a rack to keep flowers oriented upward and placed in a cooler for transport to the laboratory. For each site, 40 flower samples were collected per plant variety.

### Nectar sampling and isolation of morphotypes

Nectar was extracted from each flower within 24 h of collection from the field using microcapillary tubes (0.5 and 1-5 µL, Drummond Scientific). For each site and cultivar, 20 nectar samples were extracted at random from the 40 collected flowers (except one farm where 21 were collected), totaling 161 *V. corymbosum* and 40 *V. myrsinites* nectar samples. Some flowers contained no extractable nectar and were discarded. Nectar volume was recorded, and the sample was diluted in sterile deionized water (100 µL; ca. 1:10^2^ dilution). From this dilution, samples were further diluted 10-fold to ca. 1:10^3^-10^4^. To quantify culturable fungi, 25 µL of 10^2^ and 10^3^ dilutions were plated to YMA media with 100 mg/L chloramphenicol to inhibit bacterial growth. For bacteria, 25 µL of 10^2^ and 10^4^ dilutions were plated to sucrose supplemented R2A media (160 g/L sucrose, 18.1 g/L R2A agar) with 100 mg/L cycloheximide to inhibit fungal growth. Similar media has been used to successfully culture nectar fungi and bacteria previously (Samuni-Blank *et al*. 2014; Vannette *et al*. 2021). Sterile controls (10% of samples for each collection) were prepared in parallel by extracting 5 µL sterile synthetic nectar and processing as described.

Plated sample dilutions were incubated at 29 °C for 2-8 days until colonies formed. Plates were then photographed, and the presence of morphologically distinct colonies was recorded. For each sample, a colony from each morphotype was further cultivated to obtain pure cultures of unknown species, which were preserved in 15% glycerol at –80 °C. Colony forming units (CFUs) were determined post hoc using the collected photographs and ImageJ (Schneider, Rasband, Eliceiri 2012) to count morphologically distinct colonies. Photographs of Arcadia samples from Farm D were lost, allowing only presence/absence morphotype identification. Therefore, data from these 20 samples was excluded from community analyses but retained in analyses using presence/absence data and for comparison of nectar sugars and volumes. In 2% of samples, one or more colonies were too numerous to count, so CFUs were assigned a value of 450 as a conservative estimate of the species abundance. CFU counts per plate were transformed to CFUs per µL using the dilution factors and total nectar volume collected; CFUs per µL data was then log_10_(x+1) transformed.

### Microbial analysis

Preserved purified isolates were recultivated for sequencing and identification. In cases where preserved cultures did not grow, we attempted to extract and amplify DNA from glycerol stocks containing non-viable cells. We preserved 380 colonies, representing 56 bacterial and 41 fungal morphotypes. From these, we successfully sequenced and annotated 100 strains. In many cases, the same morphotype was repeatedly isolated from different samples within a location, so 2-3 representative isolates were sequenced to identify the strain and confirm colonies belonged to the same species.

Genomic DNA from cultured microbes was extracted using the Zyppy plasmid miniprep kit (Zymo Research, Irvine, CA) according to the manufacturer’s recommendations. Phusion High-Fidelity DNA Polymerase kits (New England BioLabs, Ipswich, MA) were used to amplify diagnostic regions of DNA for species identification. For bacteria, 16S rRNA was amplified using the 27F (AGAGTTTGATCMTGGCTCAG) and 1492R (TACGGYTACCTTGTTACGACTT) primers. For non-yeast fungi, the internal transcribed spacer region (ITS) of the ribosomal DNA was amplified using the ITS4 (GGAAGTAAAAGTCGTAACAAGG) and ITS5 (TCCTCCGCTTATTGATATGC) primers, For yeast, 28S rRNA was amplified using the NL1 (GCATATCAATAAGCGGAGGAAAAG) and NL4R (GGTCCGTGTTTCAAGACGC) primers. PCR products were visualized on agarose gels, excised, and purified using the Zymoclean Gel DNA recovery kit. Purified PCR products were Sanger sequenced at Molecular Cloning Laboratories (South San Francisco, CA) and sequences were queried against the curated 16S, 28S, or ITS databases using blastn on NCBI (blast.ncbi.nlm.nih.gov). Isolates were assigned putative identities based on highest similarity to known species and sequences were deposited in GenBank NCBI (Table S1). When a sequence matched multiple known species with similar BLAST scores, we assigned putative identity of the isolate to the genus level only. Species that were isolated only once (*n*=25) were considered likely to originate from outside the nectar samples and were removed from the dataset prior to community analysis.

### Analyses of microbial community data

Data was analysed using R Statistical Software (v4.2.2; R Core Team 2022). Linear models were fit using the **lme4** (Bates et al. 2015) and **lmerTest** (Kuznetsova, Brockhoff, Christensen 2017) packages and model performance was evaluated by examining residuals using the *check_model* function in the **easystats** package (Lüdecke et al. 2022). Statistical significance for linear model factors was evaluated using analysis of variance (ANOVA) with the *anova* function using Kenward-Roger approximation of degrees of freedom. Estimated marginal means were calculated based on the models using the *emmeans* function in the package **emmeans** (Lenth 2022).

To investigate differences in total fungal and bacterial cell densities between *Vaccinium* species, we used linear models including species as a fixed variable and location as a random variable. To compare cell densities between *V. corymbosum* samples, we first attempted to fit models with cultivar, management, and their interaction as fixed factors and a random effect of location. However, we identified significant heteroscedasticity and poor normality in the residuals of these models due to the high proportion of Arcadia samples that contained no microbes. Based on these diagnostics, we chose to instead examine cultivar and management in separate models, each with random intercepts for farm. Only Meadowlark samples were used to examine the influence of farm management.

We investigated microbial community differences with principal coordinate analysis and permutational ANOVA based on Jensen-Shannon distances using the *adonis2* function in the **vegan** package (Dixon 2003). Permutational ANOVA was used to analyse differences between cultivated *V. corymbosum* and wild *V. myrsinites* blueberries with permutations constrained by location. To identify whether microbial communities differed between organic vs. conventional management and *V. corymbosum* blueberry cultivars, we constrained the dataset to only cultivated *V. corymbosum* samples which contained microbes and used a permutational ANOVA with fixed factors of cultivar and management and permutations constrained by location. Based on these results and to further explore the influence of different farms on nectar communities, we performed another permutational ANOVA with fixed effects of location and cultivar. Homogeneity of dispersions between groups was analysed via the *betadisper* function and Tukey HSD post-hoc analyses with *p* values adjusted for multiple comparisons.

To determine which microbial species differentiate *V. corymbosum* cultivars, we used random forest models (**randomForest** package; Liaw and Wiener, 2002) and evaluated the models using out-of-bag (OOB) error rates and the area under the receiver operating characteristic curve (AUC) using the *prediction* and *performance* functions in the **ROCR** package (Sing et al. 2005).

The relationship between fungal and bacterial cell density was investigated with a linear model with response variable of bacterial cell density and fixed factor of fungal abundance with location as a random effect. To evaluate co-occurrence of fungi and bacteria within samples, we used a chi-square test of independence and a co-occurrence matrix. To identify associations between specific microbe species, we used a co-occurrence matrix which described the presence/absence data for each microbe and excluded any microbe that occurred in less than 5% of samples. We used the *cooccur* function in the **cooccur** package (Griffith, Veech, Marsh 2016) to calculate the observed and expected frequencies of co-occurrence for microbial pairs *(F_obs_* and *F_exp_*) and the probabilities that the pair would co-occur at a frequency less than or more than was observed (assuming the species were distributed randomly). These probabilities can be interpreted as *p* values, and when taken with a comparison between *F_obs_* and *F_exp_*, used to evaluate whether a pair of microbes have a positive, negative, or random association.

### Nectar chemical and data analysis

Initial dilutions of floral nectar samples described above were further diluted to ca. 1:300 with water and analysed by liquid chromatography evaporative light scattering detection as described in Rering et al. (2021). External calibration curves were used to measure fructose, glucose, and sucrose.

To analyse differences in nectar sugar concentrations between the plants without potential interference by microbial metabolism, only samples that contained no culturable microorganisms were fit with linear models including plant type (*Vaccinium myrsinites*, *V. corymbosum* cv. Arcadia, or *V. corymbosum* cv. Meadowlark) and random effect of location. Then to explore the effects of fungi and bacteria on nectar sugars, we used linear models including fungal density, bacterial density, and their interaction with random effect of location. Because sucrose was rarely detected, we evaluated the response variables fructose, glucose, and total sugars (sum of hexoses and sucrose) in our analyses. All sugar concentrations and cell densities were log-transformed.

## Results

### Microbial occurrence in blueberry nectar

Approximately 66% of samples contained culturable microbes (131 of 200 flowers).

Bacteria were detected in 62% (*n=*124) and fungi in 38.5% (*n*=77) of nectar samples. A significant positive association between the occurrence of fungi and bacteria in nectar was identified (*χ^2^*=42.5, *df*=1, *p*<0.0001), with most samples hosting culturable fungi also containing bacteria (34.5% of samples, *n*=69). Samples containing only bacteria were also common (28% of samples, *n*=56), but only six samples contained fungi and no culturable bacteria (3% of samples). Linear model results confirmed the positive association between fungi and bacteria (Fig. 1, *F_1,78_*=78.3, *p*<0.0001).

**Figure 1.**
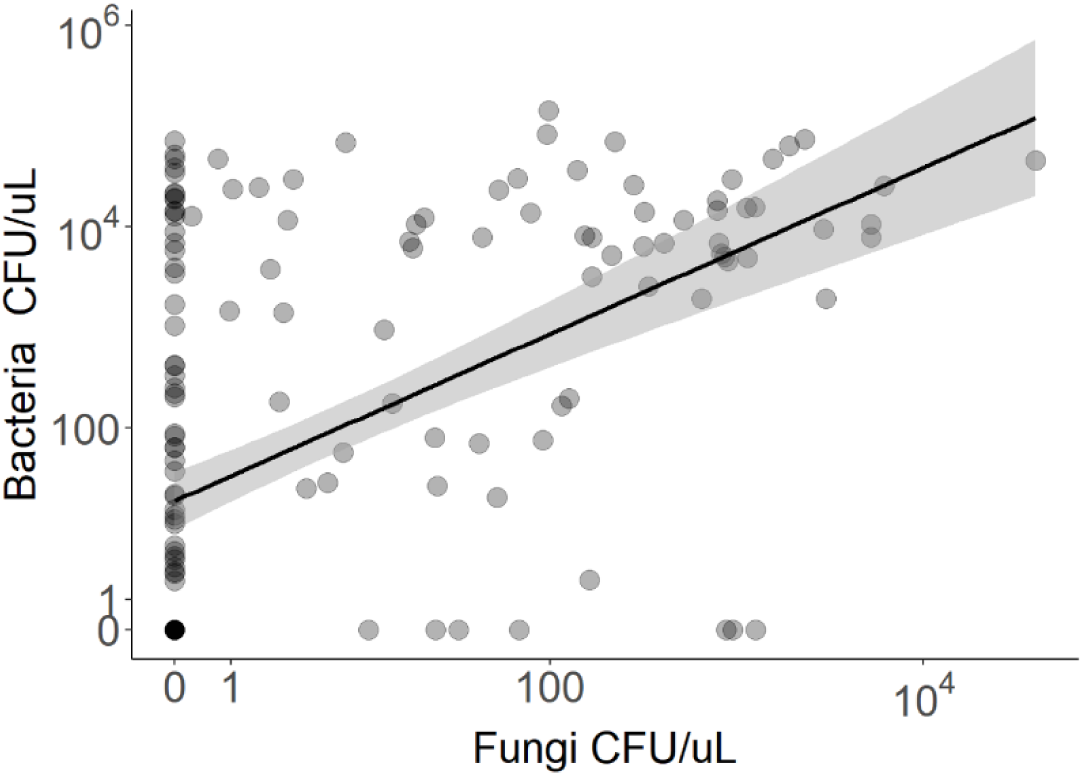
Microbial abundance in *Vaccinium* spp. nectar. Bacterial cell density is plotted as a function of fungal cell density and a linear fit is displayed with shaded region depicting the 95% confidence interval.

Thirty-two significant associations were identified between pairs of microbe species (*p<*0.05; Table S2). Of these, the majority were positive (78%, *n*=25). Positive associations were evenly split between bacteria-bacteria and bacteria-fungi pairs (13 and 12, respectively). Among negative associations, five were between bacteria-only pairs and two were between fungi-bacteria pairs. No significant associations, positive or negative, were detected between fungi-only pairs.

On average, floral nectar hosted ca. two species (0.5 fungi and 1.5 bacteria per sample). The maximum number of species isolated from a single nectar sample was seven (three fungi and four bacteria), but this unusual diversity was detected in only one flower. When microbes were present, mean cell densities were 966 and 5,952 cells per µL for fungi and bacteria, respectively. The highest cell density of a single species was achieved by a *Bacillus* sp., which occurred at 1.43 x 10^5^ cells per µL in a wild *Vaccinium myrsinites* blueberry flower collected from Ross Prairie Forest. The highest fungal density, 3.97 x 10^4^ cells per µL, was achieved by the red yeast *Symmetrospora symmetrica* in an organic Meadowlark *V. corymbosum* blueberry flower.

### Characterization of blueberry nectar communities

A total of 384 isolates were collected and preserved. Of these, 111 (71 bacteria and 40 fungi) morphologically distinct colonies from each location were cultured and sequenced. 105 isolates representing 37 species were successfully sequenced and identified to at least the genus level. Of the identified species, 16 were detected in only one sample and were removed from the dataset prior to community analysis. The remaining 21 strains included 15 bacteria and 6 fungi. An unidentified bacterium and fungus were also included in the community analysis, though their occurrence was rare (Table 1). The most frequently detected and abundant bacteria were species from the genera *Neokomagataea* (*n*=65 nectar samples)*, Rosenbergiella* (*n*=62), and *Bacillus* (*n*=38), and the most frequently occurring and abundant fungi were *Metschnikowia* (*n*=40) and *Symmetrospora* (*n*=36).

**Table 1.** Microorganisms identified in *Vaccinium* floral nectar. For each plant species and cultivar, the percentage frequency of detection and the mean cell density when isolated is provided for all microbial species (*V. corymbosum* cv. Meadowlark *n*=80, *V. corymbosum* cv. Arcadia *n*=61, *V. myrsinites n*=40).

| Kingdom | Species | *V. corymbosum* cv.<br>Meadowlark |  | *V. corymbosum* cv.<br>Arcadia |  | *V. myrsinites* |  |
| --- | --- | --- | --- | --- | --- | --- | --- |
|  |  | Mean cell<br>density<br>(CFU/μL) | Frequency<br>(%) | Mean cell<br>density<br>(CFU/μL) | Frequency<br>(%) | Mean cell<br>density<br>(CFU/μL) | Frequency<br>(%) |
| Bacteria | *Neokomagataea thailandica* | 4021 | 53.75 | 1009 | 3.28 | 0 | 0 |
|  | *Rosenbergiella* sp. | 8153 | 36.25 | 1263 | 22.95 | 4164 | 20 |
|  | *Pantoea agglomerans* | 670 | 33.75 | 133 | 1.64 | 0 | 0 |
|  | *Bacillus* sp. | 8928 | 22.5 | 3.2 | 1.64 | 30 559 | 27.5 |
|  | *Kocuria rhizophila* | 2636 | 17.5 | 0 | 0 | 0 | 0 |
|  | *Acinetobacter apis* | 2666 | 13.75 | 531 | 11.48 | 0 | 0 |
|  | *Acinetobacter nectaris* | 414 | 13.75 | 0 | 0 | 0 | 0 |
|  | *Erwinia aphidicola* | 9909 | 12.5 | 3974 | 1.64 | 0 | 0 |
|  | *Neokomagataea tanensis* | 6414 | 6.25 | 0 | 0 | 5441 | 37.5 |
|  | *Acinetobacter boissieri* | 11 981 | 5 | 3304 | 1.64 | 0 | 0 |
|  | *Lysinibacillus chungkukjangi* | 188 | 2.5 | 0 | 0 | 0 | 0 |
|  | *Acinetobacter radioresistens* | 1116 | 1.25 | 126 | 4.92 | 0 | 0 |
|  | *Pseudomonas* sp. | 1353 | 1.25 | 0 | 0 | 5552 | 17.5 |
|  | *Bacillus nealsonii* | 0 | 0 | 2452 | 4.92 | 0 | 0 |
|  | *Gluconobacter wancherniae* | 0 | 0 | 0 | 0 | 15 988 | 32.5 |
|  | unknown bacteria 1 | 0 | 0 | 9.6 | 3.28 | 0 | 0 |
| Fungi | *Symmetrospora symmetrica* | 2129 | 33.75 | 17.9 | 3.28 | 2420 | 15 |
|  | *Metschnikowia rancensis* | 187 | 25 | 0 | 0 | 10.4 | 5 |
|  | *Metschnikowia reukaufii* | 2.3 | 2.5 | 0 | 0 | 633 | 40 |
|  | *Sporidiobolus pararoseus* | 1.7 | 6.25 | 0 | 0 | 0 | 0 |
|  | *Cladosporium* sp. 1 | 0.70 | 1.3 | 0 | 0 | 9.9 | 2.5 |
|  | *Cladosporium* sp. 2 | 0.57 | 6.25 | 0 | 0 | 0 | 0 |
|  | unknown fungi 1 | 0.31 | 2.5 | 0 | 0 | 0 | 0 |

### Comparing the nectar microbiome of Vaccinium species

*V. corymbosum* and *V. myrsinites* blueberry nectar microbial communities had equal diversity (beta-dispersion *p*=0.7) and microbial cell density for both bacteria and fungi were equal in *V. myrsinites* and *V. corymbosum* samples (Fig. 2A-B; *F_1,4_*=0.076, *p*=0.8 for bacteria; *F_1,4_*=0.95, *p*=0.4 for fungi). Permutational ANOVA results did not detect differences in the microbial communities of cultivated *V. corymbosum* and wild *V. myrsinites* blueberry nectar (species *F_1,179_*=29.9, *p*=1).

**Figure 2.**
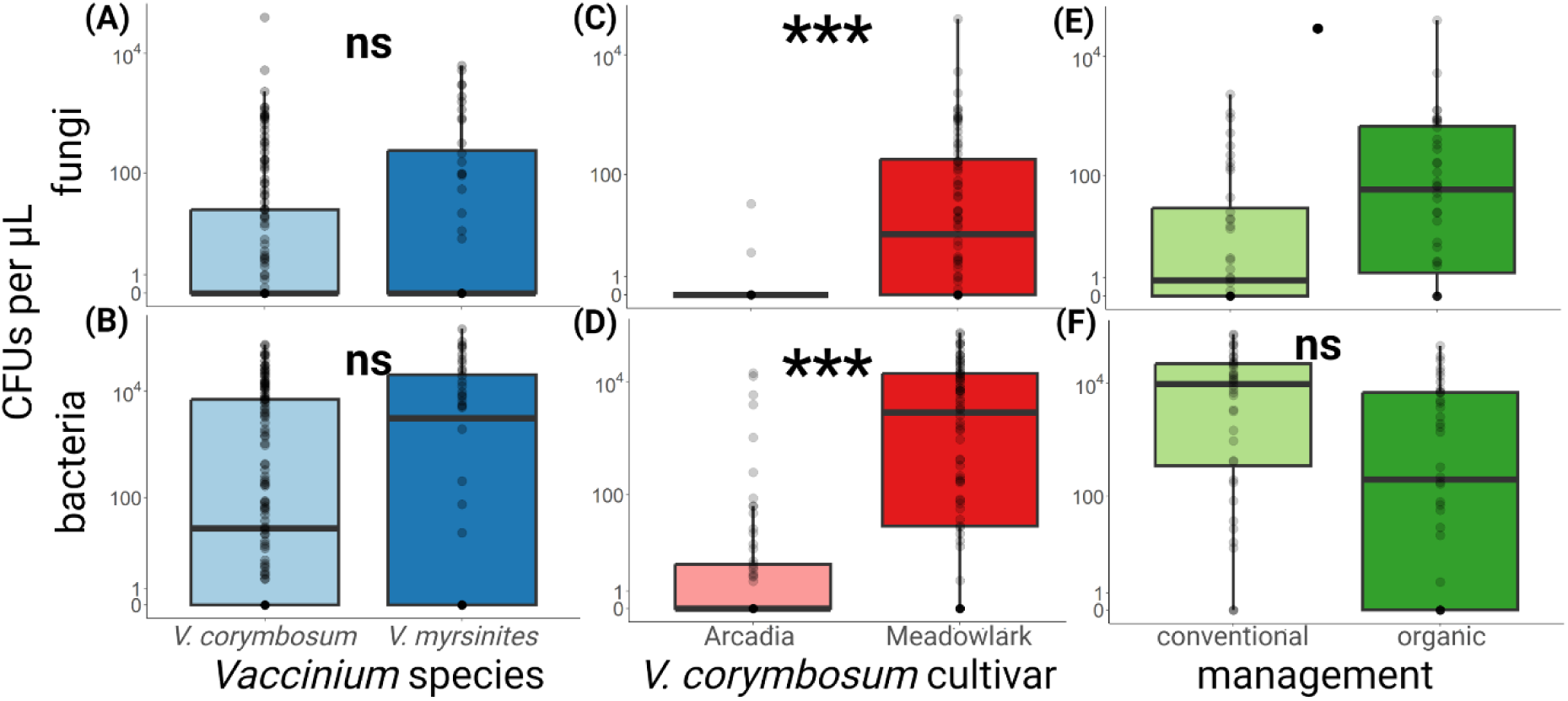
Microbial abundance in *Vaccinium corymbosum* and *V. myrsinites* nectar **(A-B)**, between *V. corymbosum* cultivars Arcadia and Meadowlark **(C-D)**, and between *V. corymbosum* cv. Meadowlark samples collected from organic or conventional farms **(E-F)** with linear model results (*** *p*<0.001, • *p*<0.1, ns = not significantly different). Boxplots depict median, first and third quartiles, and 95% confidence intervals. Individual samples are overlaid in light grey.

### Effects of farm, management, and cultivar on Vaccinium corymbosum nectar communities

Total fungal and bacterial cell density differed between cultivars, with Meadowlark containing significantly higher cell densities than Arcadia (Fig. 2C-D; bacteria *F_1,139_*=63.1, *p*<0.001; fungi *F_1,138_*=64.5, *p*<0.001). Microbial incidence was also higher in Meadowlark nectar than Arcadia, with 88% of Meadowlark samples hosting culturable microbes and only 38% of Arcadia samples. Because of the low incidence of microbes in Arcadia flowers, we evaluated the effects of management on total microbial density among Meadowlark flowers only. We detected weak evidence that organic flowers had higher fungal cell densities than conventional flowers (Fig. 2E, *F_1,2_*=8.8, *p*=0.098), but bacterial abundance was similar between crop management regimes (Fig. 2F, *F_1,2_*=2.1, *p*=0.28).

Differences in diversity were detected between farms, management regimes, and cultivars. Flowers from farm C had a relatively consistent microbiome, leading to lower group variance than was found in other farms (beta-dispersion *p*<0.001 for all comparisons with Farm C, Table S3). Nectar from the other farms, A, B, and D, had similar variance (beta-dispersion *p*≥0.44; see Table S3 for full results). Arcadia floral nectar, which often lacked culturable microbes, had lower diversity than Meadowlark (beta-dispersion *p*<0.001). Organic nectar also had lower sample variance than nectar from plants cultivated under conventional management (beta-dispersion *p*<0.001). Further investigation revealed that differences in variance between sample groups was caused by unequal distribution of microbe-free samples. When only *V. corymbosum* samples that contained microbes were analysed, sample variance between cultivars (beta-dispersion *p*=0.82), management (beta-dispersion *p*=0.21), and location were similar (beta-dispersion, *p_adj_*≥0.75, see Table S4 for full results). This reduced dataset included 52 conventional flowers (*n*=15 Arcadia, *n*=37 Meadowlark) and 41 organic flowers (*n*=8 Arcadia, *n*=33 Meadowlark).

To visualize and analyze differences between samples with culturable microbes we used PCoA (Fig. 3) and permutational ANOVA with fixed factors of cultivar, farm management and their interaction, and location as a random factor. Arcadia and Meadowlark cultivars hosted differing communities (Fig. 3A; cultivar, *F_1,89_*=11.3, *p*=0.001). Although we did not detect differences between nectar cultivated under organic vs. conventional management regimes (management, *F_1,89_*=12.3, *p*=0.64), the interaction between cultivar and management was significant (*F_1,89_*=8.87, *p*=0.001) indicating the effect of management type on community composition was different depending on cultivar.

**Figure 3.**
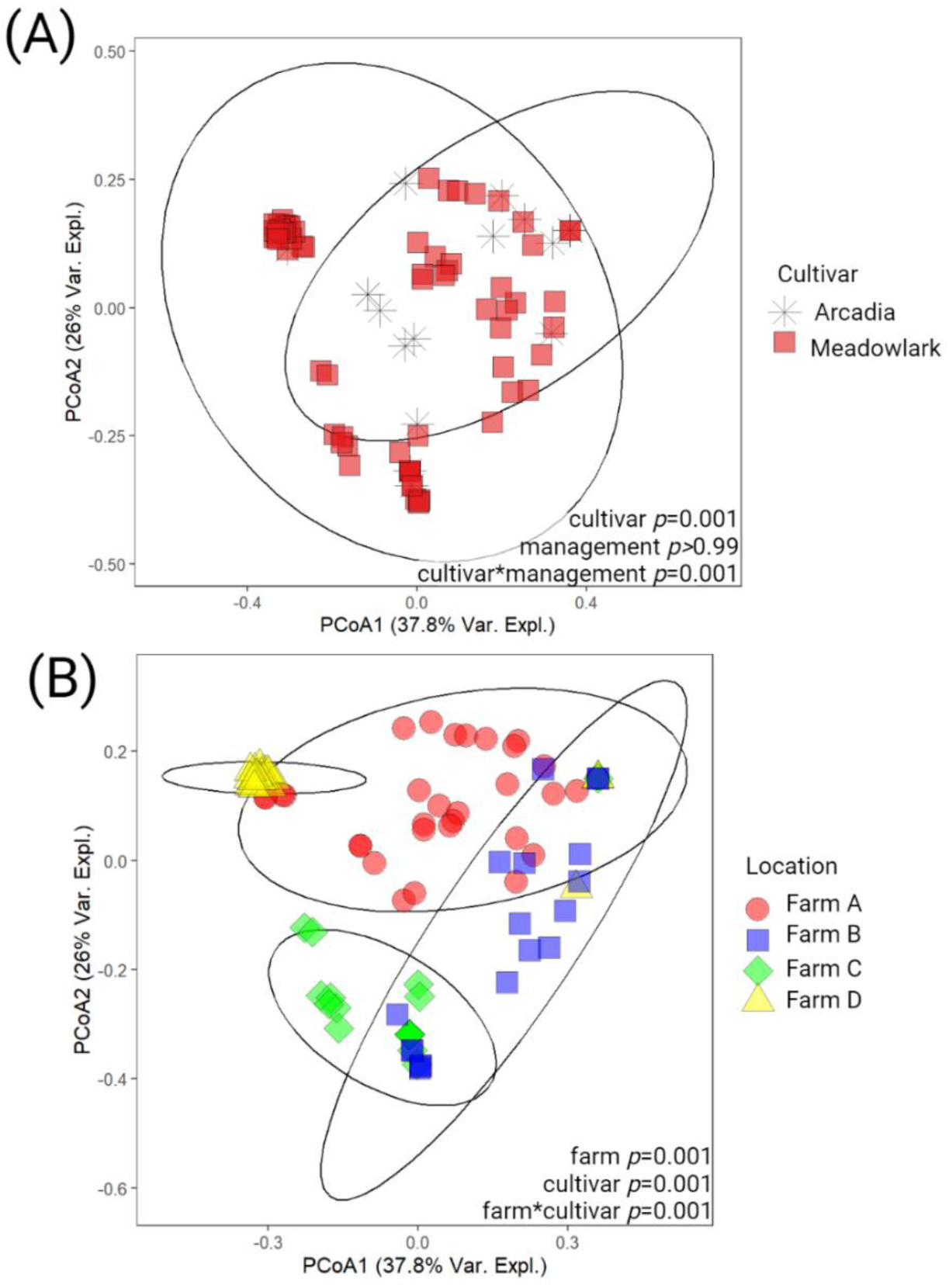
Principal coordinate analysis (PCoA) and permutational ANOVA results of Jensen-Shannon distances for nectar microbial communities of *Vaccinium corymbosum* samples that contained microbes. Two *V. corymbosum* cultivars, Arcadia and Meadowlark, were co-cultivated and collected from four farms. Permutational ANOVA results in **(A)** include cultivar and management (organic or conventional) as fixed effects and farm as a random effect. Sample colors and shape illustrate cultivars. Results in **(B)** include farm and cultivar as fixed effects and sample colors and shapes illustrate differences between farms. Ellipses indicate 95% confidence intervals for cultivar and farm in **(A)** and **(B)**, respectively.

Random forest classification models used to evaluate which microbes contribute to community differences between *V. corymbosum* cultivars ranked the bacteria *Neokomagataea thailandica* (mean decrease in Gini 5.89), *Rosenbergiella* sp. (3.41), and *Bacillus* sp. (2.97) and the yeast *Symmetrospora symmetrica* (3.58) as important in distinguishing between groups (9.7% OOB error rate, AUC 0.93; see Tables S5 and S6 for model results and variable rankings). *N. thailandica, Rosenbergiella*, *Bacillus* sp., and *S. symmetrica* were more abundant and frequent in Meadowlark than Arcadia nectar (Table 1).

Because we observed clustering of samples by farm in the PCoA (Fig. 3B), we performed an additional permutational ANOVA with fixed effects of cultivar, farm, and their interaction. Here, we confirmed that individual farms had differing communities. Farm site explained 37% of the community variance (farm, *F_3,86_*=23.2, R^2^=0.37, *p*=0.001), cultivar explained 9% variance (cultivar, *F_1,86_*=17.7, R^2^=0.09, *p*=0.001), and their interaction explained an additional 9% variance (farm*cultivar, *F_2,86_*=8.73, R^2^=0.09, *p*=0.001).

### Nectar traits differed between blueberry species and cultivars

Collected nectar volumes ranged from 0.078 to 28.711 µL per flower. Flowers that contained no extractable nectar were presumed to be recently foraged and were excluded from our sampling efforts. Nectar volume was different between the three plant varieties (Fig. 4A; *F_2,10_*=42.9, *p*<0.001). *V. corymbosum* Meadowlark blueberry flowers contained 7.5 ± 0.83 µL nectar, more than Arcadia flowers (3.4 ± 0.4 µL, *p*<0.0001) and native *V. myrsinites* blueberry flowers (0.45 ± 0.06 µL, *p*=0.06). Arcadia and *V. myrsinites* flowers had similar nectar volumes (*p*=0.3).

**Figure 4.**
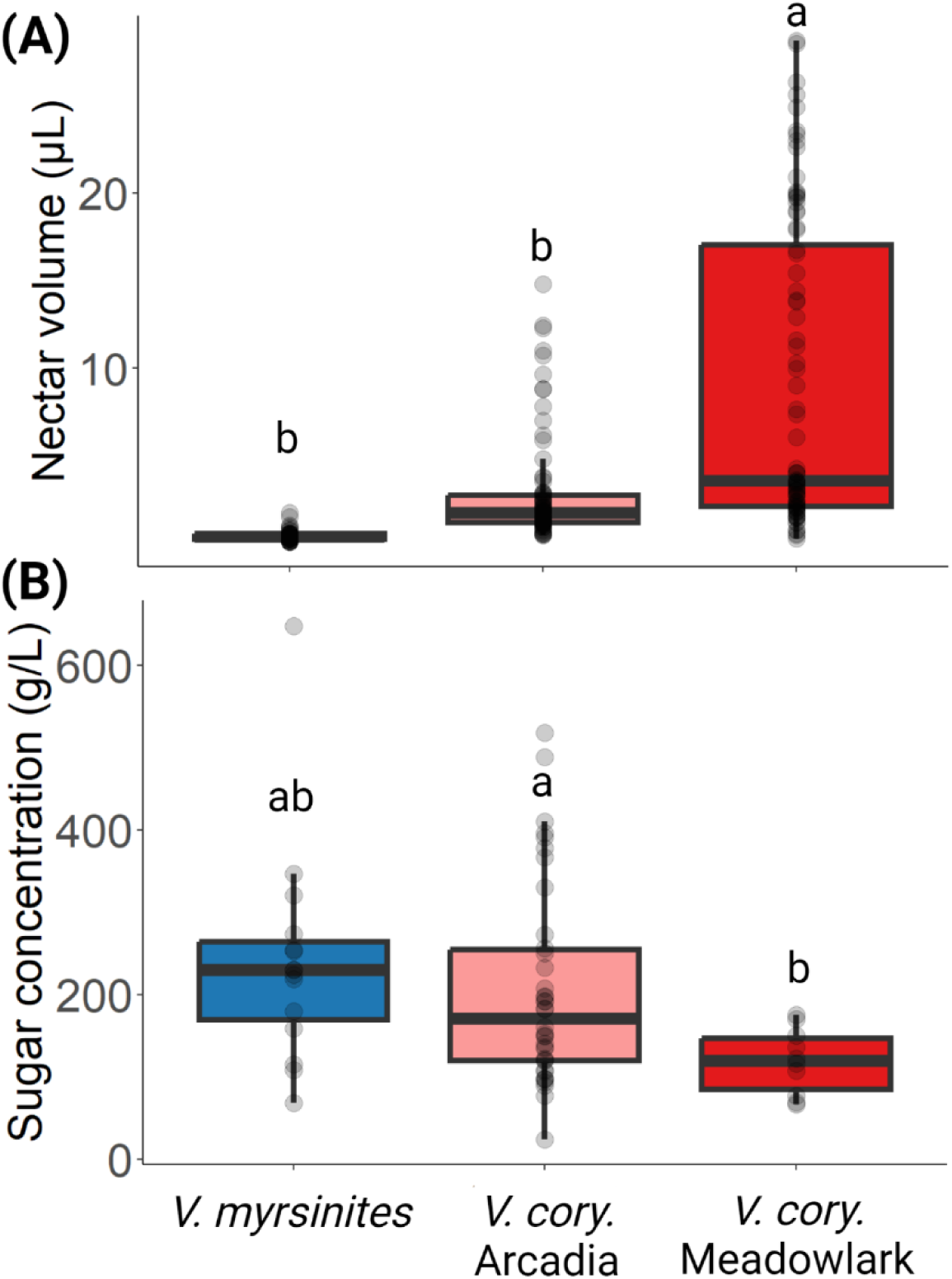
Nectar volumes **(A)** and sugar concentrations **(B)** for *Vaccinium myrsinites*, and two *V. corymbosum* cultivars, Arcadia and Meadowlark. Boxplots depict median, first and third quartiles, and 95% confidence intervals. Individual samples are overlaid in light grey. Groups labelled with different letters are different (*p*≤0.05). In **(B)**, only samples that did not contain culturable microbes were included to evaluate differences in sugar concentrations without potential effect of microbial metabolism

To investigate the carbohydrate composition of nectar as produced by the plants and without the potential interference of microbial metabolism, we constrained our dataset to only samples that did not contain culturable microorganisms. Microbe-free blueberry nectar contained abundant glucose and fructose (mean ± standard error 99 ± 8 and 87 ± 7 g/L for fructose and glucose, respectively), but sucrose was rarely detected, and when found, was at much lower concentrations (0.3 ± 0.12 g/L sucrose). Total sugar concentration differed between the plant varieties (Fig. 4B; *F_2,8_*=4.53, *p*=0.048). Meadowlark nectar (119 ± 7 g/L, 11.9% *w/v*) was less concentrated than nectar from Arcadia flowers (220 ± 30 g/L, 22% *w/v*; *p*=0.011), while *V. myrsinites* nectar (240 ± 38 g/L, 24% *w/v*) was not significantly different from either *V. corymbosum* cultivar (*V. myrsinites* vs. Meadowlark *p*=0.3, *V. myrsinites* vs. Arcadia *p*>0.9). Nectar glucose had similar differences between plants and patterns of abundance where *V. myrsinites* = Arcadia > Meadowlark (*F_2,8_*=7.73, *p*=0.012), but nectar fructose was similar in all three *Vaccinium* species (*F_2,8_*=3.02, *p*=0.11; see Fig. S2 and Table S7 for pairwise comparisons of constituent sugars between plants).

### High cell density is correlated to reduced nectar sugars

Overall, nectar sugar concentrations were negatively correlated with microbial cell density (Fig. 5A). Bacteria, fungi, and their interaction impacted total nectar sugars and glucose (*p*≤0.033 for all, see Table 2 for ANOVA results). Fructose concentrations however were strongly affected by fungi (*F_1,175_*=14.5, *p*<0.001), but only weak evidence was found for bacteria (*F_1,177_*=3.31, *p*=0.07) and the interaction between these factors (*F_1,174_*=14.5, *p*=0.07). When comparing nectar that contained no culturable microbes, fungi, bacteria, or both, microbial colonization tended to reduced sugar concentrations and lower concentrations were observed when both fungi and bacteria were present in nectar (Fig 5B).

**Figure 5.**
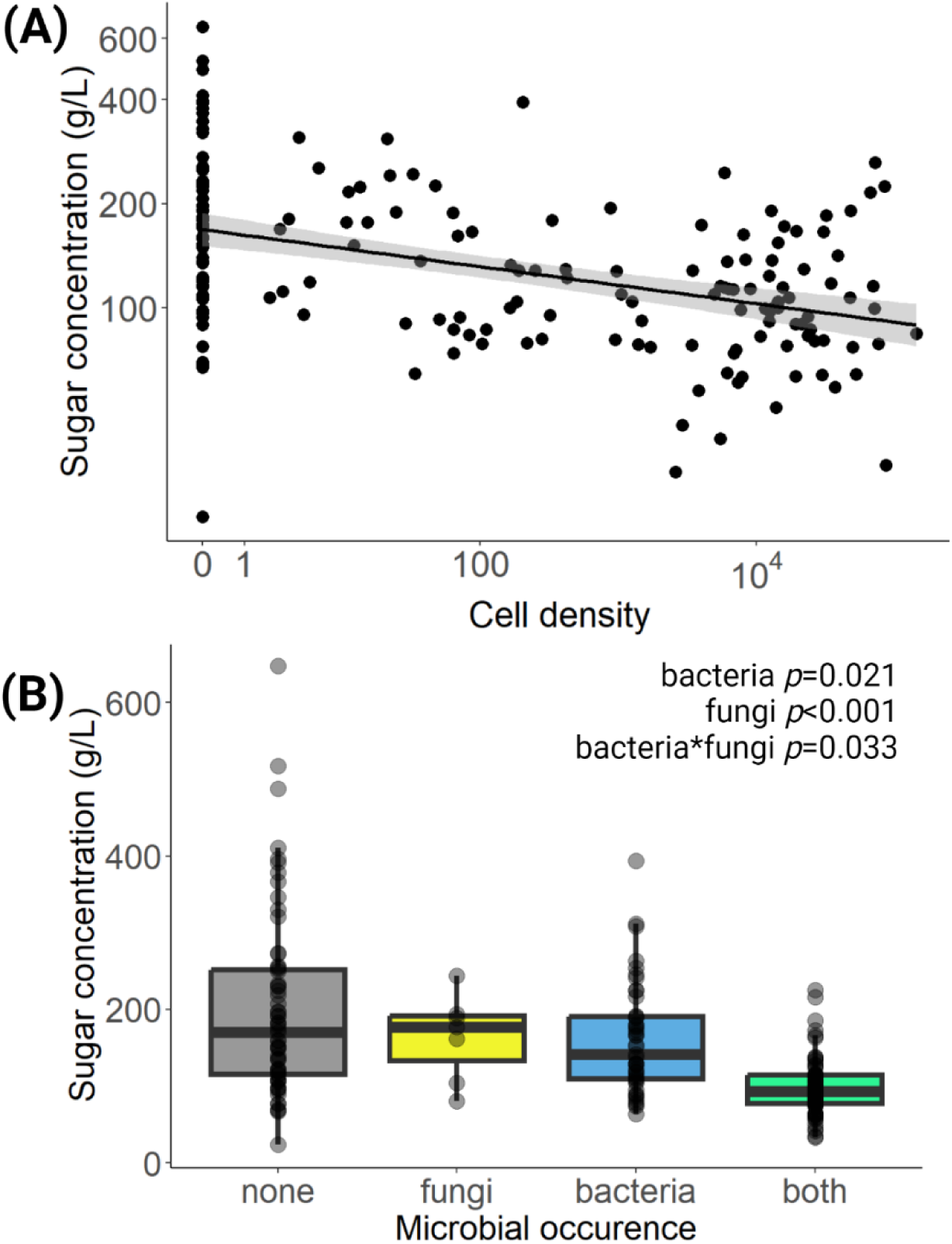
Relationship between total nectar sugar concentrations (sucrose and hexoses) and microbial cell densities. Each point represents a *Vaccinium* nectar sample. In **(A)**, cell density describes the sum of bacteria and fungi and a linear fit is displayed with shaded regions depicting the 95% confidence interval. In **(B)**, nectar sugar concentrations are displayed according to whether a sample contained no microbes, only fungi, only bacteria, or both bacteria and fungi. Linear model results for total sugars with fixed factors of fungal density, bacterial density, and their interaction with location as a random effect are displayed.

**Table 2.**
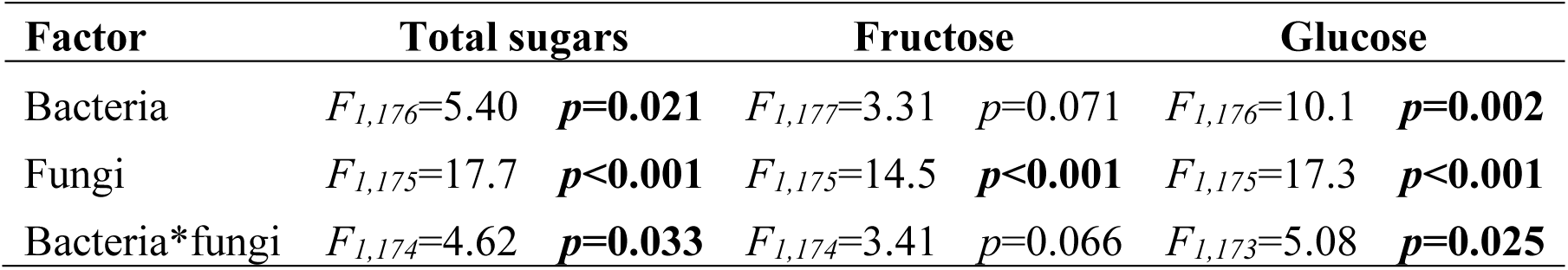
Analysis of variance results for linear models of fixed factors bacteria, fungi, and their interaction and random effect of location for dependent variables total sugars (sum of sucrose and hexoses), fructose, and glucose.

| **Factor** | **Total sugars** |  | **Fructose** |  | **Glucose** |  |
| --- | --- | --- | --- | --- | --- | --- |
| Bacteria | $F_{1,176}=5.40$ | $p=0.021$ | $F_{1,177}=3.31$ | $p=0.071$ | $F_{1,176}=10.1$ | $p=0.002$ |
| Fungi | $F_{1,175}=17.7$ | $p<0.001$ | $F_{1,175}=14.5$ | $p<0.001$ | $F_{1,175}=17.3$ | $p<0.001$ |
| Bacteria*fungi | $F_{1,174}=4.62$ | $p=0.033$ | $F_{1,174}=3.41$ | $p=0.066$ | $F_{1,173}=5.08$ | $p=0.025$ |

## Discussion

### Characterizing the Vaccinium nectar microbiome

Here we provide the first systematic exploration of blueberry nectar microbial communities, quantitatively describing the bacteria, fungi, and yeast of two crop cultivars under organic and conventional management and a native species growing in the wild. To date, only a handful of studies have simultaneously and quantitatively measured microbial abundance of both fungi and bacteria in nectar (Bartlewicz *et al*. 2016; Vannette and Fukami 2017; von Arx *et al*. 2019; Vannette *et al*. 2021; Schaeffer *et al*. 2023). These studies are needed to form a more complete conceptualization of the nectar microbiome, which is critical to plant and pollinator health and interactions.

Bacteria were more abundant and had higher incidence than fungi in blueberry nectar, as has been observed in other plants (Jacquemyn *et al*. 2013a; 2013b; Vannette *et al*. 2021), though greater frequency of yeasts compared to bacteria has also been observed (de Vega *et al*. 2021). Overall, we found bacteria and fungi co-occurred in nectar more frequently than would be anticipated by chance. Specific species were also frequently detected in the same nectar drops, including some pairs that have been previously identified as positively associated with one another, like *Acinetobacter* and *Metschnikowia* (Álvarez-Pérez and Herrera 2013).

Complementary resource utilization has been proposed as potential drivers of co-occurrence, for example preferential utilization of different nectar sugars: glucose by *Metschnikowia* and fructose by *Acinetobacter* (Álvarez-Pérez, Lievens, Fukami 2019). Co-occurrence could also be explained by a shared vector mechanism to flowers, e.g., floral visitors. We also observed negative associations between species, suggesting competition or exclusion among certain microbes.

As expected, nectar specialists such as species of *Rosenbergiella,* which has been previously isolated from blueberry flowers (Chacón et al 2022), and members of the genera *Metschnikowia*, *Acinetobacter*, and *Neokomagataea* (Álvarez-Pérez and Herrera 2013; de Vega *et al*. 2021; Vannette *et al*. 2021) were frequently detected. Additionally, insect-associated microbes were identified, like *Acinetobacter apis*, isolated from the digestive tract of honey bees (Kim et al. 2014), and *Erwinia aphidicola,* isolated from the gut of aphids (Harada et al. 1997). One somewhat unusual nectar inhabitant identified in our study was the leaf-associated yeast *Symmetrospora symmetrica* (basionym *Sporobolomyces symmetricus*, Haelewaters *et al*. 2020). Although microbes prevalent on leaves are common in nectar and floral tissues, and related *Sporobolomyces* species have been detected in nectar and flowers previously (Golonka and Vigalys 2013; Pozo, Lievens, Jacquemyn 2015; Wei *et al*. 2021), ours is the first report of *Symmetrospora symmetrica* as a major member of a plant’s nectar microbiome. This yeast was identified in cultivated and wild plants, reached the highest densities of any fungi in our study, and its frequency in floral nectar rivaled that of *Metschnikowia* species, which are regarded as quintessential nectar specialists.

Most floral microbes are culturable, and previous investigations of nectar microbial communities have identified overlap between culture-dependent and independent approaches (Morris et al. 2020; Hayes et al. 2021). However, we cannot rule out the possibility that some species were not detectable using our methods. We also caution that this survey provides data for only a single timepoint, peak bloom at each location, and we sampled a single floral stage, mature flowers exhibiting no signs of senescence. Because of this, temporal variations of nectar microbial communities remain unexplored in this system, such as variations occurring within a growing season (Schaeffer, Vannette, Irwin 2015; von Arx *et al*. 2019; Schaeffer *et al*. 2023), between years (Vannette *et al*. 2021), and as a function of flower age (Shade, McManus, Handelsman 2013; Bogo *et al*. 2021).

### Effects of location on the nectar microbiome

The blueberry nectar microbiome was highly correlated with location. The importance of site-specific factors like the local climate, land cover, flora, and fauna are known to impact microbial community assembly in flowers (Samuni-Blank *et al*. 2014; Aizenberg-Gershtein, Izhaki, Halpern 2017; Vannette *et al*. 2017; Schaeffer *et al*. 2021; Hietaranta, Juottonen, Kytöviita 2023; Schaeffer *et al*. 2023). Floral visitors are particularly significant to the establishment of nectar microbial communities. Greater visitation results in higher microbial diversity (Vannette and Fukami 2017) and density (von Arx *et al*. 2019). Furthermore, the identity of floral visitors can be more important to community composition than plant relatedness (Zemenick, Vannette, Rosenheim 2012) or geographic region (de Vega *et al*. 2021). Given that pollinator abundance and identity often vary between blueberry farms, much of the site-specific variation identified between farms in our study could be explained by differences in floral visitors. For example, all the farms in this study were stocked with honey bees, but only two (Farms A and B) were stocked with bumble bees (Mallinger, Ternest, Naranjo 2021). Wild pollinator prevalence also varies between blueberry farms and is associated with the amount of surrounding natural land (Benjamin, Reilly and Winfree 2014; Vega *et al*. 2023).

### Effects of cultivar on the nectar microbiome

Striking dissimilarities were identified between the nectar communities of *Vaccinium corymbosum* cultivars. Meadowlark nectar had higher microbial abundance and incidence than Arcadia. Even when comparing only samples where microbes were present, floral nectar communities differed between cultivars, with Meadowlark containing higher populations of several taxa including the nectar microbes *Neokomagataea thailandica* and *Rosenbergiella* sp., the honey bee gut microbe *Acinetobacter apis*, and phyllosphere-associated *Symmetrospora symmetrica*. Because both cultivars were grown at each of the farms in our study, it is likely these differences in nectar communities are mediated by cultivar-specific traits.

Arcadia and Meadowlark were selected due to differences in yields reported by Florida farmers; Meadowlark plants suffer from low yields, while Arcadia is ranked as one of the highest yielding cultivars (Phillips 2019a; 2019b). Farmers suspect that differences in yield between the cultivars are driven by differential pollinator attraction; they report high bee visits to Arcadia and low visits Meadowlark flowers (D. Phillips, pers. comm.). We did not monitor pollinator or insect visits in this study; however, given that bees prefer nectar that is very rich in sugars (Afik, Dag, Shafir 2006; Cnaani, Thosmon, Papaj 2006) and Arcadia floral nectar is ca. twice as concentrated as Meadowlark, sugar content may contribute to the increased pollinator visitation to Arcadia flowers.

If Arcadia flowers are more frequently visited by pollinators, we may expect higher microbial incidence in Arcadia nectar relative to Meadowlark as visitors introduce microbes to floral nectar. Instead, we observed the opposite, with Arcadia nectar having lower microbial incidence and abundance than Meadowlark. Assuming reports of pollinator preference are correct, there are several potential explanations for this apparent discrepancy. Since nectar cell densities increase as flowers age (von Arx *et al*. 2019), it seems most likely that differences between the cultivars are attributable to differences in the lifespan of individual flowers, e.g., Meadowlark flowers persist longer than Arcadia, either due to higher pollination requirements or innate differences between the cultivars. In addition to potential effects of flower aging, visits by non-pollinating insects like flower thrips could also contribute to Meadowlark nectar communities. Thrips can efficiently vector microbes, especially bacteria (Vannette *et al*. 2021), and blueberry flowers are often infested by these pests (Liburd *et al*. 2009).

### Effects of farm management on the nectar microbiome

In addition to differences between cultivars, we noted more subtle effects of farm management on nectar microbial communities. Organic farms had higher fungal abundance than conventional farms and farm management practices had only cultivar-specific effects on microbial community composition. Fungicides are routinely applied to blueberry blooms to prevent disease. When applied to flowers, fungicides inhibit the growth of nectar yeasts (Álvarez-Pérez et al. 2016; Bartlewicz et al. 2016) and reduce fungal richness and diversity (Schaeffer *et al*. 2017a). However, because pollinators often avoid foraging from fungicide-treated flowers and resources (Liao, Wu, Berenbaum 2017; Tamburini *et al*. 2021), they can also have indirect effects on floral microbial communities by affecting pollinator foraging (Wei *et al*. 2021).

### Comparison of Vaccinium corymbosum and V. myrsinites nectar microbial communities

In contrast to the differences noted between *Vaccinium corymbosum* cultivars, microbial abundance and community composition were similar in *V. corymbosum* and *V. myrsinites* nectar. However, we noted some interesting differences in taxa present in wild vs. cultivated flowers. For example, *Acinetobacter apis* and *Erwinia aphidicola* were only detected in cultivated flowers, suggesting a greater prevalence of honey bees and aphids in cultivated *V. corymbosum* samples and potential “imprint” of these insects on nectar communities. Also, distinct species of *Metschnikowia* were detected in cultivated vs. wild blueberry: *M. reukaufii* was much more abundant and frequent in *V. myrsinites*, whereas *M. rancensis* (basionym *Candida rancensis,* Kurtzman, Robnett, Basehoar 2018) was more common in *V. corymbosum*. Priority effects, where early arriving species suppress the growth of late comers, have been observed between these yeasts (Peay, Belisle, Fukami 2012) and can result in nectar microbes persisting in a given flower patch across multiple floral generations (Toju *et al*. 2017). This phenomenon may explain the divergent occurrence of *M. rancensis* and *M. reukaufii* between plants.

### Nectar traits differ between plants and are influenced by microbial communities

High microbial abundance was associated with reduced nectar sugars, suggesting that nectar microbes altered blueberry nectar chemistry via metabolism. Microbial metabolism of nectar sugars is recognized as a driver of nectar sugar variation in multiple other plant species (Canto and Herrera 2012; de Vega and Herrera 2013; Schaeffer, Vannette and Irwin 2015; Vannette and Fukami 2017). When microbes were absent, significant variations in nectar sugars were still evident, likely reflecting innate differences between plants. Nectar volumes also varied, although because flowers were not protected from foraging before collection, differences could to some extent reflect differential nectar removal by foragers, as opposed to innate differences in nectar production.

## Outlook

By examining blueberry crops and their wild relatives in parallel, we gained insight into the impacts of management and domestication on nectar traits and microbial communities. Interestingly, variation was apparently disconnected from genetic relatedness in this study, with the largest differences in nectar volumes, concentrations, and microbiomes found between *V. corymbosum* cultivars as opposed to between plant species. Cultivar trait selection therefore seems critical in dictating nectar microbial communities and rewards. Crop management was also a factor, with nectar from conventionally managed farms containing lower fungal abundance.

Though the potential impacts of the nectar microbes isolated in our study on blueberries are yet to be fully explored, given previous reports, they can be anticipated to have wide-ranging impacts on plants and pollinators, from negative to positive. For example, some members of *Cladosporium*, a fungus identified here, can be plant pathogenic. Conversely, other microbes may be beneficial. For example, *M. reukaufii* suppressed the growth of a floral pathogen (Crowley-Gall *et al*. 2022) and has been found in some instances to increase pollinator affinity (Vannette, Gauthier and Fukami 2013; Herrera, Pozo and Medrano 2013; Schaeffer and Irwin 2014; Schaeffer *et al*. 2017b; Yang *et al*. 2019; Colda *et al*. 2021). Moreover, although nectar bacteria seem to generally reduce pollinator attraction to nectar (Vannette, Gauthier and Fukami 2013; Good *et al*. 2014; Junker *et al*. 2014; Rering *et al*. 2018), *Acinetobacter nectaris* increased visitation by insects by honey bees and hoverflies to pear flowers in an agricultural setting (Colda *et al*. 2021). Given that blueberry yield and quality are pollination dependent, nectar microbes have the potential to improve or reduce the harvest. Therefore, increased knowledge and management of the floral microbiome could provide a sustainable method to improve crop production of pollinator dependent plants.

## Acknowledgements

We are grateful to the farmers who participated in this study by allowing us to collect flowers from their fields. We thank A. Lanier, T. Corcoran, and other student volunteers for laboratory assistance. Thanks also to D. Phillips, G. Broadhead, R. Schaeffer, R. Vannette, J. Cromie, and I.D.B. Oliveira for their helpful discussion and advice.

