## Supporting Information for "A quantitative survey of the blueberry (*Vaccinium* spp.) nectar microbiome: variation between cultivars, locations, and farm management approaches"

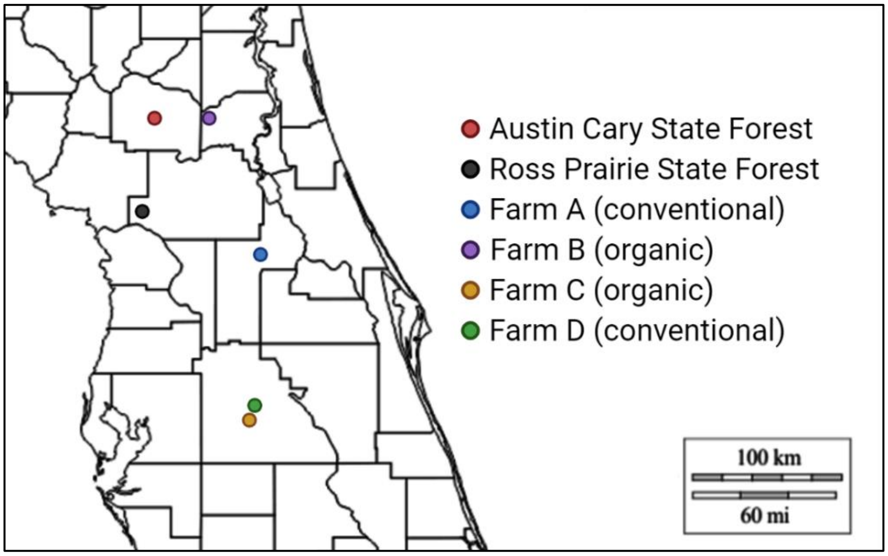


**Supplementary Figure S1.** Map of North and Central Florida. Points represent farms and state forests where blueberry flowers were collected.

**Supplementary Table S1.** Fungal and bacterial colonies isolated from floral nectar. Isolates were identified by amplifying and sequencing the 16S rRNA for bacteria, 28S rRNA for yeast, and the ITS region for non-yeast fungi. Sequences were annotated against and deposited in GenBank NCBI. N/A indicates sequencing was not successful. When an isolate matched multiple species with equal score and percent coverage, all matches were listed, and the isolate was identified to the genus level.

| Accession number | Organism | Isolate No. | Location | Plant species | Management | Date collected | Region | Primers | bp length | Top Hit | % identity | Hit Accession Number |
|---|---|---|---|---|---|---|---|---|---|---|---|---|
| OP205142 | *Metschnikowia rancensis* | 1 | Farm A | highbush | conventional | 1/30/2020 | 28S | NL1/NL4 | 514 | *Metschnikowia rancensis* | 99.80% | NG_055309.1 |
| N/A | N/A | 2 | Farm A | highbush | conventional | 1/30/2020 | N/A | N/A | N/A | N/A | N/A | N/A |
| OP205143 | *Sporidiobolus pararoseus* | 3 | Farm D | highbush | conventional | 2/19/2020 | 28S | NL1/NL4 | 588 | *Sporidiobolus pararoseus* | 99.66% | NG_067256.1 |
| OP205144 | *Moesziomyces aphidis* | 4 | Farm D | highbush | conventional | 2/19/2020 | 28S | NL1/NL4 | 626 | *Moesziomyces aphidis* | 100.00% | NG_069796.1 |
| OP205145 | *Metschnikowia reukaufii* | 5 | Farm C | highbush | organic | 2/19/2020 | 28S | NL1/NL4 | 453 | *Metschnikowia reukaufii* | 98.90% | NG_055410.1 |
| OP205146 | *Sporidiobolus pararoseus* | 6 | Farm D | highbush | conventional | 2/19/2020 | 28S | NL1/NL4 | 589 | *Sporidiobolus pararoseus* | 99.66% | NG_067256.1 |
| OP205147 | *Metschnikowia rancensis* | 7 | Farm B | highbush | organic | 2/17/2020 | 28S | NL1/NL4 | 520 | *Metschnikowia rancensis* | 99.80% | NG_055309.1 |
| OP595808 | *Sarocladium strictum* | 8 | Farm D | highbush | conventional | 2/19/2020 | ITS | ITS4/ITS5 | 581 | *Sarocladium strictum* | 98.78% | NR_111145.1 |
| OP595634 | *Methylobacterium tardum* | 11 | Farm C | highbush | organic | 2/12/2020 | 16S | 27F/1492R | 1338 | *Methylobacterium tardum* | 99.85% | NR_041443.1 |
| OP205148 | *Metschnikowia reukaufii* | 12 | Austin Cary | shiny | wild | 3/3/2020 | 28S | NL1/NL4 | 456 | *Metschnikowia reukaufii* | 99.33% | NG_055410.1 |
| OP205149 | *Metschnikowia peoriensis* | 13 | Austin Cary | shiny | wild | 3/3/2020 | 28S | NL1/NL4 | 524 | *Metschnikowia peoriensis* | 100.00% | NG_064474.1 |
| OP205150 | *Metschnikowia reukaufii* | 14 | Ross Prairie | shiny | wild | 3/3/2020 | 28S | NL1/NL4 | 317 | *Metschnikowia reukaufii* | 99.68% | NG_055410.1 |
| OP205151 | *Metschnikowia rancensis* | 15 | Ross Prairie | shiny | wild | 3/3/2020 | 28S | NL1/NL4 | 524 | *Metschnikowia rancensis* | 99.80% | NG_055309.1 |
| OP595809 | *Cladosporium* sp. | 16 | Farm D | highbush | conventional | 2/19/2020 | ITS | ITS4/ITS5 | 544 | *Cladosporium oxysporum/ Cladosporium tenuissimum* | 100.00% | NR_152267.1/NR_119855.1 |
| OP595810 | *Brunneomyces brunnescens* | 17 | Farm D | highbush | conventional | 2/19/2020 | ITS | ITS4/ITS5 | 579 | *Brunneomyces brunnescens* | 99.06% | NR_145408.1 |
| OP595811 | *Alternaria alstroemeriae* | 18 | Farm D | highbush | conventional | 2/19/2020 | ITS | ITS4/ITS5 | 569 | *Alternaria alstroemeriae* | 99.82% | NR_163686.1 |
| N/A | N/A | 19 | Farm D | highbush | conventional | 2/19/2020 | ITS | ITS4/ITS5 | 605 | *Ophiostoma acarorum* | 78.56% | NR_158859.1 |
| OP595635 | *Fructobacillus tropaeoli* | 20 | Farm B | highbush | organic | 2/10/2020 | 16S | 27F/1492R | 671 | *Fructobacillus tropaeoli* | 100.00% | NR_113043.1 |
| OP595636 | *Bombella apis* | 21 | Farm B | highbush | organic | 2/10/2020 | 16S | 27F/1492R | 1350 | *Bombella apis* | 99.63% | NR_157653.1 |
| N/A | N/A | 22 | Farm B | highbush | organic | 2/10/2020 | N/A | N/A | N/A | N/A | N/A | N/A |
| OP595637 | *Leuconostoc mesenteroides* | 23 | Farm B | highbush | organic | 2/10/2020 | 16S | 27F/1492R | 648 | *Leuconostoc mesenteroides* | 99.69% | NR_074957.1 |
| OP595638 | *Neokomagataea thailandica* | 24 | Farm A | highbush | conventional | 1/30/2020 | 16S | 27F/1492R | 556 | *Neokomagataea thailandica* | 99.13% | NR_112958.1 |
| OP595639 | *Neokomagataea thailandica* | 25 | Farm A | highbush | conventional | 1/30/2020 | 16S | 27F/1492R | 471 | *Neokomagataea thailandica* | 99.57% | NR_112958.1 |
| OP595640 | *Rosenbergiella* sp. | 26 | Farm A | highbush | conventional | 1/30/2020 | 16S | 27F/1492R | 531 | *Rosenbergiella australiborealis/ Rosenbergiella collisarenosi/ Rosenbergiella epipactidis/ Rosenbergiella nectarea* | 100.00% | NR_126305.1/NR_126304.1/NR_126303.1/NR_117969.1 |

| | | | | | | | | | | | | |
|---|---|---|---|---|---|---|---|---|---|---|---|---|
| OP595641 | *Pantoea agglomerans* | 27 | Farm A | highbush | conventional | 1/30/2020 | 16S | 27F/1492R | 666 | *Pantoea agglomerans* | 99.85% | NR_114111.1 |
| OP595642 | *Pantoea agglomerans* | 28 | Farm A | highbush | conventional | 1/30/2020 | 16S | 27F/1492R | 868 | *Pantoea agglomerans* | 99.08% | NR_114111.1 |
| OP595643 | *Fructobacillus fructosus* | 29 | Farm B | highbush | organic | 2/10/2020 | 16S | 27F/1492R | 603 | *Fructobacillus fructosus* | 100.00% | NR_113579.1 |
| OP205152 | *Symmetrospora symmetrica* | 30 | Farm A | highbush | conventional | 1/30/2020 | 28S | NL1/NL4 | 615 | *Symmetrospora symmetrica* | 99.51% | NG_057632.1 |
| OP205153 | *Symmetrospora symmetrica* | 31 | Farm A | highbush | conventional | 1/30/2020 | 28S | NL1/NL4 | 610 | *Symmetrospora symmetrica* | 99.67% | NG_057632.1 |
| OP205154 | *Symmetrospora symmetrica* | 32 | Farm D | highbush | conventional | 2/19/2020 | 28S | NL1/NL4 | 608 | *Symmetrospora symmetrica* | 99.67% | NG_057632.1 |
| OP205155 | *Symmetrospora symmetrica* | 34 | Ross Prairie | shiny | wild | 3/3/2020 | 28S | NL1/NL4 | 604 | *Symmetrospora symmetrica* | 99.67% | NG_057632.1 |
| OP205156 | *Symmetrospora symmetrica* | 35 | Austin Cary | shiny | wild | 3/3/2020 | 28S | NL1/NL4 | 606 | *Symmetrospora symmetrica* | 99.67% | NG_057632.1 |
| OP205157 | *Symmetrospora symmetrica* | 36 | Farm B | highbush | organic | 2/10/2020 | 28S | NL1/NL4 | 576 | *Symmetrospora symmetrica* | 99.65% | NG_057632.1 |
| OP205158 | *Symmetrospora symmetrica* | 37 | Farm B | highbush | organic | 2/10/2020 | 28S | NL1/NL4 | 609 | *Symmetrospora symmetrica* | 99.51% | NG_057632.1 |
| OP595812 | *Zasmidium nocoxi* | 38 | Farm C | highbush | organic | 2/12/2020 | ITS | ITS4/ITS5 | 537 | *Zasmidium nocoxi* | 99.26% | NR_156536.1 |
| OP595813 | *Staninwardia suttonii* | 39 | Farm C | highbush | organic | 2/12/2020 | ITS | ITS4/ITS5 | 559 | *Staninwardia suttonii* | 94.23% | NR_137118.1 |
| OP595644 | *Acinetobacter radioresistens* | 40 | Farm A | highbush | conventional | 1/30/2020 | 16S | 27F/1492R | 1347 | *Acinetobacter radioresistens* | 100.00% | NR_114074.1 |
| OP595645 | *Bacillus nealsonii* | 41 | Farm A | highbush | conventional | 1/30/2020 | 16S | 27F/1492R | 1367 | *Bacillus nealsonii* | 99.73% | NR_044546.1 |
| OP595646 | *Rosenbergiella* sp. | 42 | Farm A | highbush | conventional | 1/30/2020 | 16S | 27F/1492R | 322 | *Rosenbergiella australiborealis/ Rosenbergiella epipactidis* | 99.69% | NR_126305.1/NR_126303.1 |
| N/A | N/A | 43 | Farm A | highbush | conventional | 1/30/2020 | 16S | N/A | N/A | N/A | N/A | N/A |
| OP205159 | *Symmetrospora symmetrica* | 45 | Farm C | highbush | organic | 2/19/2020 | 28S | NL1/NL4 | 580 | *Symmetrospora symmetrica* | 99.66% | NG_057632.1 |
| OP595647 | *Neokomagataea thailandica* | 46 | Farm A | highbush | conventional | 1/30/2020 | 16S | 27F/1492R | 1357 | *Neokomagataea thailandica* | 99.63% | NR_112958.1 |
| OP595648 | *Neokomagataea thailandica* | 47 | Farm A | highbush | conventional | 1/30/2020 | 16S | 27F/1492R | 1357 | *Neokomagataea thailandica* | 99.63% | NR_112958.1 |
| OP595649 | *Rosenbergiella epipactidis* | 48 | Farm A | highbush | conventional | 1/30/2020 | 16S | 27F/1492R | 1426 | *Rosenbergiella epipactidis* | 99.86% | NR_126303.1 |
| OP595650 | *Acinetobacter apis* | 49 | Farm A | highbush | conventional | 1/30/2020 | 16S | 27F/1492R | 1416 | *Acinetobacter apis* | 99.93% | NR_133952.1 |
| OP595651 | *Acinetobacter apis* | 50 | Farm A | highbush | conventional | 1/30/2020 | 16S | 27F/1429R | 1419 | *Acinetobacter apis* | 99.93% | NR_133952.1 |
| OP595652 | *Acinetobacter apis* | 51 | Farm A | highbush | conventional | 1/30/2020 | 16S | 27F/1492R | 1417 | *Acinetobacter apis* | 99.93% | NR_133952.1 |
| OP595653 | *Acinetobacter apis* | 52 | Farm A | highbush | conventional | 1/30/2020 | 16S | 27F/1492R | 1410 | *Acinetobacter apis* | 99.93% | NR_133952.1 |
| OP595654 | *Micrococcus yunnanensis* | 53 | Farm A | highbush | conventional | 1/30/2020 | 16S | 27F/1492R | 1389 | *Micrococcus yunnanensis* | 99.64% | NR_116578.1 |
| OP595655 | *Acinetobacter apis* | 54 | Farm A | highbush | conventional | 1/30/2020 | 16S | 27F/1492R | 1409 | *Acinetobacter apis* | 99.93% | NR_133952.1 |
| OP595656 | *Kocuria rhizophila* | 55 | Farm B | highbush | organic | 2/17/2020 | 16S | 27F/1492R | 443 | *Kocuria rhizophila* | 99.77% | NR_026452.1 |
| OP595657 | *Acinetobacter nectaris* | 56 | Farm B | highbush | organic | 2/17/2020 | 16S | 27F/1492R | 1411 | *Acinetobacter nectaris* | 99.29% | NR_118408.1 |
| OP595658 | *Acinetobacter apis* | 57 | Farm B | highbush | organic | 2/17/2020 | 16S | 27F/1492R | 1412 | *Acinetobacter apis* | 99.93% | NR_133952.1 |
| OP595659 | *Neokomagataea thailandica* | 58 | Farm B | highbush | organic | 2/17/2020 | 16S | 27F/1492R | 1354 | *Neokomagataea thailandica* | 99.63% | NR_112958.1 |
| OP595660 | *Neokomagataea thailandica* | 59 | Farm B | highbush | organic | 2/17/2020 | 16S | 27F/1492R | 1361 | *Neokomagataea thailandica* | 99.63% | NR_112958.1 |
| OP595661 | *Pantoea agglomerans* | 60 | Farm B | highbush | organic | 2/17/2020 | 16S | 27F/1492R | 1409 | *Pantoea agglomerans* | 99.36% | NR_041978.1 |
| OP595662 | *Pantoea agglomerans* | 61 | Farm B | highbush | organic | 2/17/2020 | 16S | 27F/1492R | 1410 | *Pantoea agglomerans* | 99.36% | NR_041978.1 |
| OP595663 | *Pantoea agglomerans* | 63 | Farm B | highbush | organic | 2/17/2020 | 16S | 27F/1492R | 1406 | *Pantoea agglomerans* | 99.43% | NR_041978.1 |
| OP595664 | *Dermacoccus nishinomiyaensis* | 64 | Farm C | highbush | organic | 2/12/2020 | 16S | 27F/1492R | 1375 | *Dermacoccus nishinomiyaensis* | 99.78% | NR_044872.1 |
| OP595665 | *Bacillus* sp. | 65 | Farm C | highbush | organic | 2/12/2020 | 16S | 27F/1492R | 1421 | *Bacillus velezensis/ Bacillus amyloliquefaciens* | 99.79% | NR_116240.1/NR_041455.1 |

| | | | | | | | | | | | | |
|---|---|---|---|---|---|---|---|---|---|---|---|---|
| OP595666 | *Rosenbergiella epipactidis* | 66 | Farm C | highbush | organic | 2/12/2020 | 16S | 27F/1492R | 1394 | *Rosenbergiella epipactidis* | 100.00% | NR_126303.1 |
| OP595667 | *Erwinia aphidicola* | 67 | Farm C | highbush | organic | 2/12/2020 | 16S | 27F/1492R | 1406 | *Erwinia aphidicola* | 100.00% | NR_104724.1 |
| OP595668 | *Rosenbergiella epipactidis* | 68 | Farm D | highbush | conventional | 2/19/2020 | 16S | 27F/1492R | 1413 | *Rosenbergiella epipactidis* | 99.65% | NR_126303.1 |
| OP595669 | *Rosenbergiella epipactidis* | 69 | Farm D | highbush | conventional | 2/19/2020 | 16S | 27F/1492R | 1388 | *Rosenbergiella epipactidis* | 99.93% | NR_126303.1 |
| OP595670 | *Pseudomonas* sp. | 70 | Farm D | highbush | conventional | 2/19/2020 | 16S | 27F/1492R | 1391 | *Pseudomonas oryzihabitans/ Pseudomonas psychrotolerans* | 99.93% | NR_114041.1/ NR_042191.1 |
| OP595671 | *Rosenbergiella epipactidis* | 71 | Farm D | highbush | conventional | 2/19/2020 | 16S | 27F/1492R | 1409 | *Rosenbergiella epipactidis* | 99.93% | NR_126303.1 |
| OP595672 | *Erwinia aphidicola* | 73 | Farm D | highbush | conventional | 2/19/2020 | 16S | 27F/1492R | 481 | *Erwinia aphidicola* | 100.00% | NR_104724.1 |
| OP595673 | *Pantoea agglomerans* | 74 | Farm D | highbush | conventional | 2/19/2020 | 16S | 27F/1492R | 1407 | *Pantoea agglomerans* | 99.57% | NR_041978.1 |
| OP595674 | *Pantoea agglomerans* | 75 | Farm D | highbush | conventional | 2/19/2020 | 16S | 27F/1492R | 985 | *Pantoea agglomerans* | 99.70% | NR_041978.1 |
| OP595675 | *Neokomagataea tanensis* | 76 | Farm C | highbush | organic | 2/19/2020 | 16S | 27F/1492R | 1354 | *Neokomagataea tanensis* | 99.48% | NR_112959.1 |
| OP595676 | *Neokomagataea thailandica* | 77 | Farm C | highbush | organic | 2/19/2020 | 16S | 27F/1492R | 1353 | *Neokomagataea thailandica* | 99.56% | NR_112958.1 |
| OP595677 | *Pseudomonas* sp. | 78 | Ross Prairie | shiny | wild | 3/3/2020 | 16S | 27F/1492R | 839 | *Pseudomonas oryzihabitans/ Pseudomonas psychrotolerans* | 99.64% | NR_114041.1/ NR_042191.1 |
| OP595678 | *Bacillus velezensis* | 79 | Ross Prairie | shiny | wild | 3/3/2020 | 16S | 27F/1492R | 1396 | *Bacillus velezensis* | 99.93% | NR_116240.1 |
| OP595679 | *Gluconobacter wancherniae* | 80 | Ross Prairie | shiny | wild | 3/3/2020 | 16S | 27F/1492R | 1351 | *Gluconobacter wancherniae* | 99.33% | NR_112952.1 |
| OP595680 | *Rosenbergiella epipactidis* | 81 | Ross Prairie | shiny | wild | 3/3/2020 | 16S | 27F/1492R | 1401 | *Rosenbergiella epipactidis* | 99.86% | NR_126303.1 |
| OP595681 | *Gluconobacter wancherniae* | 82 | Ross Prairie | shiny | wild | 3/3/2020 | 16S | 27F/1492R | 1352 | *Gluconobacter wancherniae* | 99.26% | NR_112952.1 |
| OP595682 | *Pantoea anthophila* | 83 | Ross Prairie | shiny | wild | 3/3/2020 | 16S | 27F/1492R | 833 | *Pantoea anthophila* | 99.76% | NR_116749.1 |
| N/A | N/A | 84 | Ross Prairie | shiny | wild | 3/3/2020 | 16S | 27F/1492R | N/A | N/A | N/A | N/A |
| OP595683 | *Gluconobacter wancherniae* | 85 | Austin Cary | shiny | wild | 3/3/2020 | 16S | 27F/1492R | 540 | *Gluconobacter wancherniae* | 99.82% | NR_112952.1 |
| OP595684 | *Neokomagataea tanensis* | 86 | Ross Prairie | shiny | wild | 3/3/2020 | 16S | 27F/1492R | 1357 | *Neokomagataea tanensis* | 99.56% | NR_112959.1 |
| OP595685 | *Lysinibacillus chungkukjangi* | 87 | Farm B | highbush | organic | 2/17/2020 | 16S | 27F/1492R | 905 | *Lysinibacillus chungkukjangi* | 99.88% | NR_109669.1 |
| OP595686 | *Rosenbergiella* sp. | 88 | Austin Cary | shiny | wild | 3/3/2020 | 16S | 27F/1492R | 1408 | *Rosenbergiella epipactidis/ Rosenbergiella collisarenosi* | 99.72% | NR_126303.1/ NR_126304.1 |
| OP595687 | *Neokomagataea thailandica* | 90 | Farm B | highbush | organic | 2/17/2020 | 16S | 27F/1492R | 1345 | *Neokomagataea thailandica* | 99.55% | NR_112958.1 |
| OP595688 | *Acinetobacter nectaris* | 91 | Farm C | highbush | organic | 2/19/2020 | 16S | 27F/1492R | 1336 | *Acinetobacter nectaris* | 98.88% | NR_118408.1 |
| OP595689 | *Neokomagataea thailandica* | 92 | Farm C | highbush | organic | 2/19/2020 | 16S | 27F/1492R | 1259 | *Neokomagataea thailandica* | 99.44% | NR_112958.1 |
| OP595690 | *Neokomagataea thailandica* | 93 | Farm A | highbush | conventional | 1/30/2020 | 16S | 27F/1492R | 1346 | *Neokomagataea thailandica* | 99.55% | NR_112958.1 |
| OP595691 | *Acinetobacter boissieri* | 94 | Farm A | highbush | conventional | 1/30/2020 | 16S | 27F/1492R | 1343 | *Acinetobacter boissieri* | 98.44% | NR_118409.1 |
| N/A | N/A | 95 | Farm A | highbush | conventional | 1/30/2020 | N/A | N/A | N/A | N/A | N/A | N/A |
| OP595692 | *Acinetobacter boissieri* | 96 | Farm B | highbush | organic | 2/17/2020 | 16S | 27F/1492R | 1356 | *Acinetobacter boissieri* | 98.82% | NR_118409.1 |
| OP595693 | *Acinetobacter boissieri* | 97 | Farm B | highbush | organic | 2/17/2020 | 16S | 27F/1492R | 1317 | *Acinetobacter boissieri* | 98.56% | NR_118409.1 |
| OP595694 | *Neokomagataea thailandica* | 98 | Farm B | highbush | organic | 2/17/2020 | 16S | 27F/1492R | 1223 | *Neokomagataea thailandica* | 99.59% | NR_112958.1 |
| OP595814 | *Papiliotrema rajasthanensis* | 99 | Farm C | highbush | organic | 2/19/2020 | ITS | ITS4/ITS5 | 530 | *Papiliotrema rajasthanensis* | 96.62% | NR_155678.1 |
| OP205160 | *Symmetrospora symmetrica* | 100 | Farm C | highbush | organic | 2/12/2020 | 28S | NL1/NL4 | 612 | *Symmetrospora symmetrica* | 99.67% | NG_057632.1 |
| OP205161 | *Metschnikowia rancensis* | 101 | Farm D | highbush | conventional | 2/19/2020 | 28S | NL1/NL4 | 525 | *Metschnikowia rancensis* | 99.80% | NG_055309.1 |
| OP595815 | *Talaromyces dendriticus* | 102 | Austin Cary | shiny | wild | 3/3/2020 | ITS | ITS4/ITS5 | 611 | *Talaromyces dendriticus* | 93.62% | NR_167968.1 |

| | | | | | | | | | | | | |
|---|---|---|---|---|---|---|---|---|---|---|---|---|
| OP595816 | *Cladosporium* sp. | 103 | Austin Cary | shiny | wild | 3/3/2020 | ITS | ITS4/ITS5 | 545 | *Cladosporium pini-ponderosae/ Cladosporium colombiae* | 99.82% | NR_119730.1/ NR_119729.1 |
| OP205162 | *Symmetrospora symmetrica* | 104 | Farm C | highbush | organic | 2/19/2020 | 28S | NL1/NL4 | 615 | *Symmetrospora symmetrica* | 99.67% | NG_057632.1 |
| OP205163 | *Symmetrospora symmetrica* | 105 | Farm C | highbush | organic | 2/19/2020 | 28S | NL1/NL4 | 612 | *Symmetrospora symmetrica* | 99.67% | NG_057632.1 |
| OP595817 | *Cladosporium* sp. | 107 | Farm C | highbush | organic | 2/19/2020 | ITS | ITS4/ITS5 | 539 | *Cladosporium pini-ponderosae/ Cladosporium colombiae* | 99.26% | NR_119730.1/ NR_119729.1 |
| OP595818 | *Cladosporium* sp. | 108 | Farm D | highbush | conventional | 2/19/2020 | ITS | ITS4/ITS5 | 542 | *Cladosporium pini-ponderosae/ Cladosporium colombiae* | 99.82% | NR_119730.1/ NR_119729.1 |
| OP205164 | *Symmetrospora symmetrica* | 109 | Farm D | highbush | conventional | 2/19/2020 | 28S | NL1/NL4 | 608 | *Symmetrospora symmetrica* | 99.67% | NG_057632.1 |
| OP205165 | *Symmetrospora symmetrica* | 110 | Farm D | highbush | conventional | 2/19/2020 | 28S | NL1/NL4 | 610 | *Symmetrospora symmetrica* | 99.51% | NG_057632.1 |
| OP595695 | *Erwinia aphidicola* | 111 | Farm D | highbush | conventional | 2/19/2020 | 16S | 27F/1492R | 1400 | *Erwinia aphidicola* | 99.86% | NR_104724.1 |

**Supplementary Table S2.** Results from a probabilistic model of microbial species co-occurrence among *Vaccinium* nectar samples. Observed incidences of each species alone and when co-isolated are reported. Calculated expected frequencies of co-occurrence and associated probability values, which can be interpreted as *p* values, are listed. Species pairs have a negative association (signified with red downward pointing arrow) when the probability of co-occurrence below what was observed is <0.05. If the probability of co-occurrence greater than what was observed is <0.05, species have a positive association (signified with green upward pointing arrow). Species pairs sorted by valence of co-occurrence probability.

| **Species 1** | **Species 2** | **species 1 incidence** | **species 2 incidence** | **Observed co-occurrence** | **Probability that both species occur at a site** | **Expected co-occurrence** | **Probability of co-occurrence less than was observed** | **Probability of co-occurrence greater than was observed** | **Species pair more or less likely to cooccur** | **Species Interaction** |
|---|---|---|---|---|---|---|---|---|---|---|
| *Bacillus* sp. | *Erwinia aphidicola* | 35 | 15 | 9 | 0.013 | 2.6 | 0.99999 | **0.00013** | ↑ | bacteria-bacteria |
| *Gluconobacter wancherniae* | *Metschnikowia reukaufii* | 13 | 18 | 10 | 0.006 | 1.2 | 1 | **0** | ↑ | fungi-bacteria |
| *Kocuria rhizophila* | *Acinetobacter apis* | 14 | 18 | 7 | 0.006 | 1.3 | 1 | **0.00003** | ↑ | bacteria-bacteria |
| *Metschnikowia rancensis* | *Acinetobacter nectaris* | 22 | 11 | 9 | 0.006 | 1.2 | 1 | **0** | ↑ | fungi-bacteria |
| *Metschnikowia rancensis* | *Kocuria rhizophila* | 22 | 14 | 12 | 0.008 | 1.5 | 1 | **0** | ↑ | fungi-bacteria |
| *Metschnikowia rancensis* | *Acinetobacter apis* | 22 | 18 | 9 | 0.01 | 2 | 1 | **0.00001** | ↑ | fungi-bacteria |
| *Neokomagataea tanensis* | *Metschnikowia reukaufii* | 20 | 18 | 14 | 0.009 | 1.8 | 1 | **0** | ↑ | fungi-bacteria |
| *Neokomagataea tanensis* | *Gluconobacter wancherniae* | 20 | 13 | 7 | 0.006 | 1.3 | 1 | **0.00004** | ↑ | bacteria-bacteria |
| *Neokomagataea tanensis* | *Bacillus* sp. | 20 | 35 | 10 | 0.017 | 3.5 | 0.99994 | **0.0004** | ↑ | bacteria-bacteria |
| *Neokomagataea thailandica* | *Acinetobacter nectaris* | 45 | 11 | 11 | 0.012 | 2.5 | 1 | **0** | ↑ | bacteria-bacteria |
| *Neokomagataea thailandica* | *Kocuria rhizophila* | 45 | 14 | 14 | 0.016 | 3.1 | 1 | **0** | ↑ | bacteria-bacteria |
| *Neokomagataea thailandica* | *Metschnikowia rancensis* | 45 | 22 | 19 | 0.025 | 4.9 | 1 | **0** | ↑ | fungi-bacteria |
| *Neokomagataea thailandica* | *Acinetobacter apis* | 45 | 18 | 12 | 0.02 | 4 | 1 | **0.00003** | ↑ | bacteria-bacteria |
| *Pantoea agglomerans* | *Kocuria rhizophila* | 33 | 14 | 12 | 0.011 | 2.3 | 1 | **0** | ↑ | bacteria-bacteria |
| *Pantoea agglomerans* | *Neokomagataea thailandica* | 33 | 45 | 19 | 0.037 | 7.4 | 1 | **0** | ↑ | bacteria-bacteria |
| *Pantoea agglomerans* | *Metschnikowia rancensis* | 33 | 22 | 12 | 0.018 | 3.6 | 1 | **0.00001** | ↑ | fungi-bacteria |
| *Pantoea agglomerans* | *Acinetobacter apis* | 33 | 18 | 8 | 0.015 | 3 | 0.99949 | **0.0031** | ↑ | bacteria-bacteria |
| *Pantoea agglomerans* | *Acinetobacter nectaris* | 33 | 11 | 6 | 0.009 | 1.8 | 0.99964 | **0.0032** | ↑ | bacteria-bacteria |
| *Pantoea agglomerans* | *Rosenbergiella* sp. | 33 | 62 | 17 | 0.051 | 10.2 | 0.9983 | **0.0056** | ↑ | bacteria-bacteria |
| *Rosenbergiella* sp. | *Pseudomonas* sp. | 62 | 10 | 6 | 0.015 | 3.1 | 0.98939 | **0.049** | ↑ | bacteria-bacteria |

| | | | | | | | | | | |
|---|---|---|---|---|---|---|---|---|---|---|
| *Symmetrospora symmetrica* | *Neokomagataea tanensis* | 36 | 20 | 10 | 0.018 | 3.6 | 0.99992 | **0.00052** | ↑ | fungi-bacteria |
| *Symmetrospora symmetrica* | *Neokomagataea thailandica* | 36 | 45 | 15 | 0.04 | 8.1 | 0.99909 | **0.0033** | ↑ | fungi-bacteria |
| *Symmetrospora symmetrica* | *Pantoea agglomerans* | 36 | 33 | 12 | 0.029 | 5.9 | 0.99887 | **0.0044** | ↑ | fungi-bacteria |
| *Symmetrospora symmetrica* | Rosenbergiella sp. | 36 | 62 | 18 | 0.055 | 11.1 | 0.99792 | **0.0065** | ↑ | fungi-bacteria |
| *Symmetrospora symmetrica* | *Bacillus* sp. | 36 | 35 | 11 | 0.031 | 6.3 | 0.99205 | **0.024** | ↑ | fungi-bacteria |
| *Bacillus* sp. | *Acinetobacter apis* | 35 | 18 | 0 | 0.016 | 3.1 | **0.027** | 1 | ↓ | bacteria-bacteria |
| *Kocuria rhizophila* | *Rosenbergiella* sp. | 14 | 62 | 0 | 0.021 | 4.3 | **0.0046** | 1 | ↓ | bacteria-bacteria |
| *Metschnikowia rancensis* | *Bacillus* sp. | 22 | 35 | 0 | 0.019 | 3.8 | **0.011** | 1 | ↓ | fungi-bacteria |
| *Neokomagataea thailandica* | *Gluconobacter wancherniae* | 45 | 13 | 0 | 0.014 | 2.9 | **0.033** | 1 | ↓ | bacteria-bacteria |
| *Pantoea agglomerans* | *Neokomagataea tanensis* | 33 | 20 | 0 | 0.016 | 3.3 | **0.023** | 1 | ↓ | bacteria-bacteria |
| *Pantoea agglomerans* | *Metschnikowia reukaufii* | 33 | 18 | 0 | 0.015 | 3 | **0.034** | 1 | ↓ | fungi-bacteria |
| *Rosenbergiella* sp. | *Acinetobacter nectaris* | 62 | 11 | 0 | 0.017 | 3.4 | **0.015** | 1 | ↓ | bacteria-bacteria |

**Supplementary Table S3.** Tukey HSD multiple comparisons of multivariate dispersion homogeneity between cultivated *V. corymbosum* samples. Mean distance between respective group medians and 95% family-wise confidence intervals with *p* values adjusted for multiple comparisons are reported. Bold *p* values indicate sample variance is statistically different between farms.

| **Group comparison** | **Difference between means** | **Lower 95% CI** | **Upper 95% CI** | ***P*-value** |
|---|---|---|---|---|
| Farm B-Farm A | -0.046 | -0.126 | 0.034 | 0.44 |
| Farm C-Farm A | -0.245 | -0.325 | -0.165 | < **0.001** |
| Farm D-Farm A | -0.009 | -0.107 | 0.090 | 1.00 |
| Farm C-Farm B | -0.199 | -0.279 | -0.118 | < **0.001** |
| Farm D-Farm B | 0.038 | -0.061 | 0.136 | 0.76 |
| Farm D-Farm C | 0.236 | 0.137 | 0.335 | < **0.001** |

**Supplementary Table S4.** Tukey HSD multiple comparisons of multivariate dispersion homogeneity between *Vaccinium corymbosum* samples that contained microbes. Mean distance between respective group medians and 95% family-wise confidence intervals with *p* values adjusted for multiple comparisons are reported.

| **Group comparison** | **Difference between means** | **Lower 95% CI** | **Upper 95% CI** | ***P*-value** |
|---|---|---|---|---|
| Farm B-Farm A | -0.047 | -0.172 | 0.077 | 0.75 |
| Farm C-Farm A | -0.003 | -0.138 | 0.0132 | 1.00 |
| Farm D-Farm A | -0.026 | -0.161 | 0.108 | 0.96 |
| Farm C-Farm B | 0.044 | -0.101 | 0.190 | 0.86 |
| Farm D-Farm B | 0.021 | -0.124 | 0.167 | 0.98 |
| Farm D-Farm C | -0.023 | -0.177 | 0.131 | 0.98 |

**Supplementary Table S5.** Random forest classification model confusion matrix results for *Vaccinium corymbosum* cultivars. The model had a 9.7% out of bag estimate of error rate. Matrix depicts distribution of samples from each group according to their predicted vs. actual classification. The class error describes the proportion of samples that were incorrectly identified within a group.

| Predicted vs. actual class | Arcadia | Meadowlark | Class error |
|---|---|---|---|
| Arcadia | 18 | 5 | 0.217 |
| Meadowlark | 4 | 66 | 0.057 |

**Supplementary Table S6.** Variable rankings for microbes in random forest classification model of *V. corymbosum* cultivars reported in S7. Higher mean decrease in Gini indicates greater importance of the microbe as a variable differentiating between the cultivars.

| **Mean decrease in Gini** | **Microbe species** |
|---:|---|
| 5.89 | *Neokomagataea thailandica* |
| 3.58 | *Symmetrospora symmetrica* |
| 3.41 | *Rosenbergiella* sp. |
| 2.97 | *Bacillus* sp. |
| 1.87 | *Acinetobacter apis* |
| 1.74 | *Pantoea agglomerans* |
| 1.06 | *Metschnikowia rancensis* |
| 1.06 | *Bacillus nealsonii* |
| 1.03 | *Acinetobacter radioresistens* |
| 0.77 | *Erwinia aphidicola* |
| 0.63 | unknown bacteria 1 |
| 0.52 | *Kocuria rhizophila* |
| 0.42 | *Cladosporium* sp. 2 |
| 0.37 | *Acinetobacter boissieri* |
| 0.34 | *Sporidiobolus pararoseus* |
| 0.26 | *Acinetobacter nectaris* |
| 0.24 | *Metschnikowia reukaufii* |
| 0.10 | *Neokomagataea tanensis* |
| 0.05 | unknown fungi 1 |
| 0.04 | *Cladosporium* sp. 1 |
| 0.02 | *Pseudomonas* sp. |
| 0.01 | *Lysinibacillus chungkukjangi* |
| 0.00 | *Gluconobacter wancherniae* |

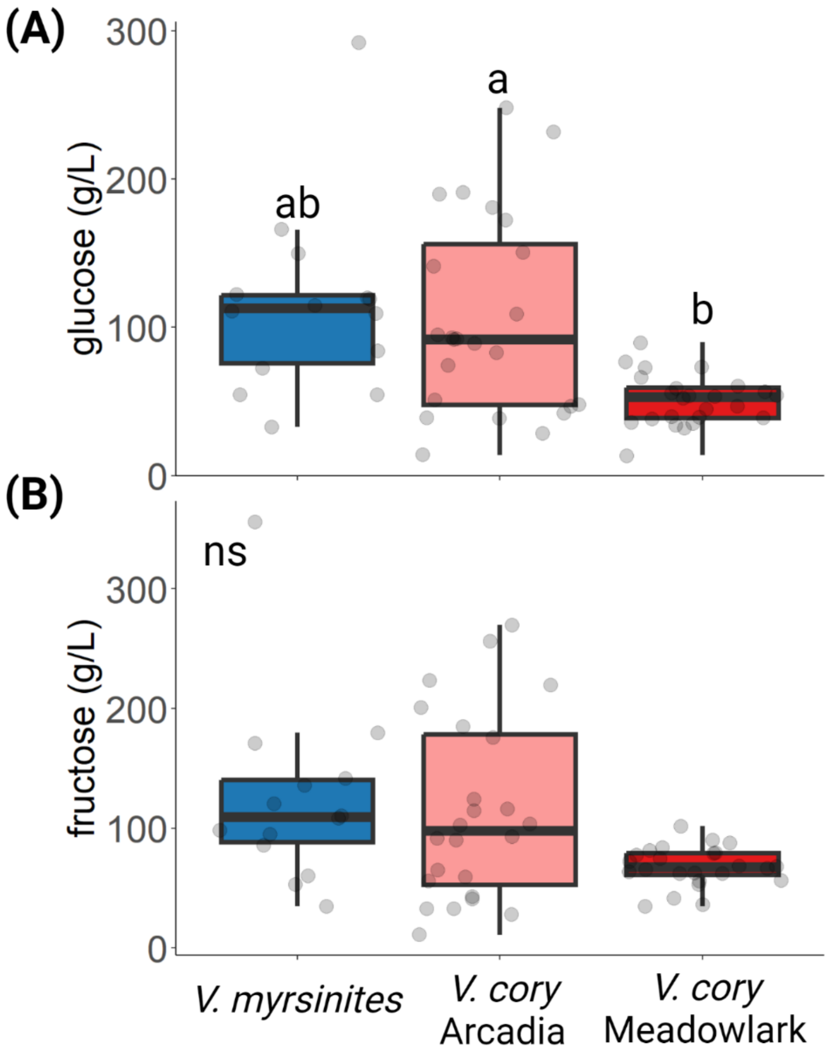


**Supplementary Figure S2.** Concentrations of A) glucose and B) fructose in blueberry nectar samples that did not contain culturable microbes. Letters denote differences between groups according to estimated marginal means from linear models ($p \leq 0.05$). *V.* myrsinites = *Vaccinium myrsinites, V. cory* = *V. corymbosum.*

**Supplementary Table S7.** Pairwise comparisons of nectar sugars between plants using estimated marginal means from linear models, *p* values adjusted for multiple comparisons. Meadowlark and Arcadia are *Vaccinium corymbosum* cultivars, shiny is the common name of *V. myrsinites*.

| Plants | *p* value | |
|---|---|---|
| | total sugars | glucose |
| Meadowlark - Arcadia | **0.011** | **0.0006** |
| Meadowlark - shiny | 0.32 | 0.21 |
| Arcadia - shiny | 0.95 | 0.94 |
